# Improvement of association between confidence and accuracy after integration of separate evidence over time

**DOI:** 10.1101/2021.06.20.449145

**Authors:** Zahra Azizi, Sajjad Zabbah, Azra Jahanitabesh, Reza Ebrahimpour

## Abstract

When making decisions in real-life, we may receive discrete evidence during a time period. Although participants can integrate information from separate cues to improve their accuracy, it is still debatable how confidence changes after receiving discrete information. Nevertheless, based on the strong positive relationship between accuracy and confidence, we predicted that similar to what is observed in accuracy, confidence would improve following the integration of separate pieces of information. We used a Random-dot-motion discrimination task in which one or two brief stimuli (i.e., pulse[s]) were presented, and participants had to indicate the predominant direction of dot motions by saccadic eye movement. Two pulses intervals (up to 1s) were randomly selected, where color-coded targets facilitated indicating confidence simultaneously. Using behavioral data, computational models, pupillometry, and EEG methodology, our data revealed that compared to single-pulse trials, in double-pulse trials, participants improve their confidence resolution rather than reporting higher confidence and that the EEG signal (centro-parietal-positivity) and pupil dynamics appear to follow confidence levels but only in double-pulse trials. Overall, our findings suggest that evidence integration helps to improve the associations between confidence and accuracy.

## 1 Introduction

To make everyday decisions, both humans and animals can integrate multiple separate pieces of information they receive from their environment. For example, imagine that a large bus is passing the street between you and a car moving at a distance. In this situation, you can easily decide whether the distant car is moving toward or away from you by integrating the separate information that can be seen through the bus windows. In this scenario, the more information you collect, the better your inference of the direction of the distant vehicle will be. Indeed, research has shown that the accuracy of decisions can be significantly improved by integrating information from separate cues (Kiani, Churchland, & Shadlen, 2013; Kira, Yang, & Shadlen, 2015; tickle, Tsetsos, Speekenbrink, & Summerfield, 2020; Tohidi-Moghaddam, Zabbah, Olianezhad, & Ebrahimpour, 2019; Waskom & Kiani, 2018).

In addition to selecting a choice among some alternatives, our decisions are accompanied by confidence —a feeling that reflects the likelihood that the decision is correct (Kiani, Corthell, & Shadlen, 2014). For example, imagine that the scene in the previous scenario also includes foggy weather. In this case, low visibility may reduce the confidence of your judgments. This diminished confidence per se may lead to change your mind (Fleming, Putten, & Daw, 2018; Resulaj, Kiani, Wolpert, & Shadlen, 2009), impact your behavioral adjustments, and affect how quickly and accurately you make your consecutive decisions (Meyniel, Sigman, & Mainen, 2015; van den Berg, Zylberberg, Kiani, Shadlen, & Wolpert, 2016). Due to the potential effects of confidence on decision-making, in the last few years, considerable progress has been made to understand of the behavioral (Kiani et al., 2014; Zylberberg, Barttfeld, & Sigman, 2012; Zylberberg, Fetsch, & Shadlen, 2016) and the neuronal (Baranski et al., 2017; Gherman & Philiastides, 2015; Kiani & Shadlen, 2009) properties of confidence and its relation with perceptual decision-making. However, it is still unclear how confidence is established if time intervals separate the provided information.

It has been shown that when we need to decide based on the separate pieces of information, the accuracy of the decision is determined by integrating the information of pieces and exceeding expectations predicted by models (Kiani et al., 2013). Furthermore, performance did not depend on the interval between the cues (Kiani et al., 2013; Waskom & Kiani, 2018), and that the second cue had the larger leverage on decisions (Kiani et al., 2013; Tohidi-Moghaddam, Zabbah, Olianezhad, & Ebrahimpour, 2019). Accordingly, one may suggest that confidence would follow the same characteristics as accuracy: increased considerably after receiving separate pieces of information, not being affected by the time interval between the cues and be leveraged larger by the most recent pack of information.

Nevertheless, a large body of evidence (e.g., [Herce Castañón et al., 2019; Zylberberg et al., 2016]) determines that human observers do not consistently report their confidence with a similar pattern of their accuracy. From this standpoint, noise can be considered as the key parameter to clarify variations in confidence (Kiani et al., 2014; Zylberberg et al., 2012). For instance, an underestimation of sensory noise in decisions would lead to over and/or under-confidence (De Gardelle & Mamassian, 2015; Herce Castañón et al., 2019; Zylberberg, Roelfsema, & Sigman, 2014). Moreover, confidence ratings may not be originated solely from the available sensory evidence (Rahnev & Denison, 2018; Zylberberg et al., 2016). For example, the observers may integrate secondary evidence into their confidence rating, which may not have been used when making the decision— utilizing secondary evidence allows the observers to change their mind after the initiating of a response (Atiya et al., 2020; Resulaj et al., 2009).

To test the hypothetical relation between accuracy and confidence, in binary decisions, signal detection theory (SDT) can provide a method to characterize the observers’ reliability in their confidence ratings by introducing metacognitive sensitivity and efficiency (Figure 1B; [Fleming, 2017; Maniscalco & Lau, 2012, 2014]). In fact, for many years, SDT has been used as a simple yet powerful methodology to distinguish between an observer’s ability to categorize the stimulus and the behavioral response (Green & Swets, 1966) and to determine confidence resolution (Balsdon, Wyart, & Mamassian, 2020; Maniscalco & Lau, 2012, 2014).

**Figure 1.**
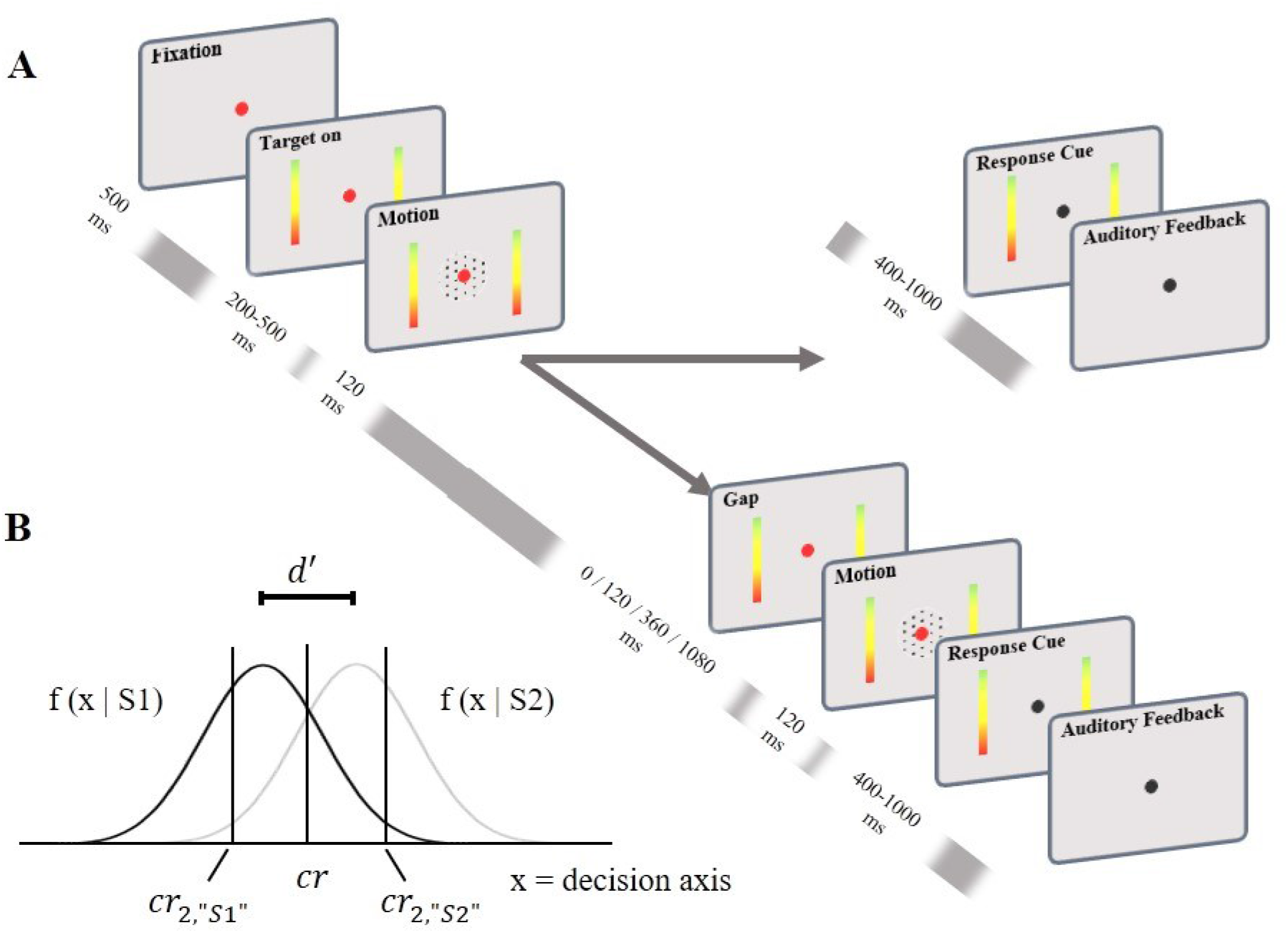
Task paradigm and Signal Detection Theory. **(A)** Participants had to indicate the predominant direction of motion of moving dots (left or right) by saccadic eye movement to one of the targets after receiving one or two pulse(s) of 120ms stimulus. The intervals between two pulses were selected randomly from 0 to 1080 ms and the direction of both pulses were the same. Color-coded targets enabled participants indicating their confidence simultaneously. **(B)** On each trial, a stimulus generates an internal response *x* within an observer, who must use *x* to decide whether the stimulus is *S*_1_ or *S*_2_, *x* is drawn from a normal distribution. The distance between these distributions is *d*′, which measures the observer’s ability to discriminate *S*_1_ from *S*_2_. The observer also rates decision confidence on a scale of high and low by comparing *x* to the additional response specific confidence criteria (*cr*_2_ for each option). For details, see Supplementary Appendix 2 and refs (Fleming, 2017; Maniscalco & Lau, 2012, 2014).

Additionally, levels of confidence can be traced by behavioral, neural, and pupillometry signatures. For example, higher confidence is accompanied by faster and more accurate decisions (Kiani et al., 2014; van den Berg et al., 2016; Zylberberg et al., 2016). Moreover, motion stimulus fluctuations influence both the accuracy (Kiani et al., 2008; Resulaj et al., 2009) and confidence (Van Den Berg et al., 2016; Zylberberg et al., 2012) and inform us about the parts of the stimulus that bear more intensely on the choice and confidence (Kiani et al., 2013, 2008; Nienborg & Cumming, 2009). Furthermore, perceptual decision-making literature has introduced an EEG potential characterized by a centro-parietal positivity (CPP) as a neural correlate of sensory evidence accumulation (Kelly & O’Connell, 2013; O’connell, Dockree, & Kelly, 2012) and confidence (Boldt, Schiffer, Waszak, & Yeung, 2019; Herding, Ludwig, von Lautz, Spitzer, & Blankenburg, 2019; Tagliabue et al., 2019; Vafaei Shooshtari et al., 2019; Zizlsperger, Sauvigny, Händel, & Haarmeier, 2014). In particular, despite the differences between correct and incorrect decisions, CPP presents regardless of whether the decision is correct and incorrect (O’connell et al., 2012; Steinemann, O’Connell, & Kelly, 2018) and can reflect not only external evidence but also an internal decision quantity such as decision confidence. Along with CPP, levels of confidence can be traced by monitoring the pupil. The literature has suggested strong links between pupil response and both the decision (Murphy, Boonstra, & Nieuwenhuis, 2016) and confidence (Allen et al., 2016; Lempert, Chen, & Fleming, 2015; Urai, Braun, & Donner, 2017) via pupil-linked dynamics of the noradrenergic system (Laeng, Sirois, & Gredebäck, 2012). For example, pupillometry has provided some evidence that shows a partial dissociation between choice and confidence in decision-making (Balsdon et al., 2020). Therefore, considering the potential of response-time, CPP, and pupillometry signatures in capturing the distinction between accuracy and confidence, they can be considered informative paradigms to explore the confidence-accuracy association in decision-making.

To bridge the existing gap in confidence and decision-making when information integrated from temporally separated stimulus compared to when only one stimulus interval is provided, we implemented two experiments to explore, as the primary goal, whether, after receiving two separate pieces of evidence, confidence reflects decision accuracy. In addition, as the secondary goals, we examined whether confidence (1) depends on the time interval between the pieces, (2) is leveraged by the second piece, and (3) predicts implicit markers—response-time, CPP and pupillometry after receiving two separate pieces of evidence. To the best of our knowledge, how evidence accumulation processes improve the accuracy and confidence association using the combination of behavioral, neural, and pupillometry signatures was not examined before.

To clarify confidence, we required observers to make a two-alternative decision after viewing either one motion pulse (single-pulse) or two motion pulses (double-pulse) separated by four various temporal gaps (similar to [Kiani et al., 2013; Tohidi-Moghaddam et al., 2019]). Considering four temporal gaps allowed us to examine whether the participants’ confidence report would vary if they respond immediately or up to 1s after the first pulse. We performed linear and logistic regression models to measure the impact of stimulus characteristics on choice and confidence. Also, we applied a set of computational models based on SDT to assess whether confidence can be predicted only based on decision information. We proposed that to predict confidence we need both decision and confidence information. Accordingly, first, we fitted the standard signal detection model to participants’ choices and confidence to directly estimate each participant’s confidence, stimulus sensitivity, and metacognitive sensitivity. Second, to explore our proposition based on the task’s nature, a perfect integrator model and two optimized models were fitted based on choices and confidence of single-pulse trials to predict confidence of double-pulse trials. Then, in the second experiment, we used EEG methodology to examine the relation between participants’ brain activity and their confidence. We expected that a neural indicator of perceptual decision making (CPP) would show amplitude changes between the two levels of confidence. Participants’ pupil response was also monitored across both experiments to examine the relation between participants’ pupil response and their confidence. Considering the evidence that the accuracy of the decision is enhanced by integrating the information over time (Kiani et al., 2013), we hypothesized that confidence would follow the same characteristics as accuracy such that participants would integrate information from pulses invariant to the temporal gap to improve their confidence. In addition, based on the previous evidence that participants obtained more information from a second pulse (Kiani et al., 2013; Tohidi-Moghaddam et al., 2019), we predicted that confidence would be leveraged larger by more recent information. Furthermore, as the effectiveness of capturing the distinction between choice and confidence has been shown using implicit markers (e.g., for response-time, see Vafaei Shooshtari et al., 2019); for CPP, see Steinemann et al., 2018; for pupillometry, see Balsdon et al., 2020), we expected that the behavioral, neural, and pupillometry markers of confidence to be distinguishable on both single-pulse and double-pulse trials.

## 2 Materials and Methods

### 2.1 Participants

Consistent with methodological considerations in previous studies, a total of 19 observers participated in two separate studies. Six participants (two males; age range [27–38], M = 32.25, SD = 4.5) attended in our behavioral study —Experiment 1— and 13 participants (three males; age range [21–39], M = 31.41; SD = 5.56) took part in our EEG study —Experiment 2. All participants had a normal or corrected-to-normal vision, and they had no history of psychiatric and neurological disorders. Previous studies with similar paradigms in which a large number of trials were presented to a small number of participants (e.g., five participants in [Kiani et al., 2013]; six participants in [Kiani et al., 2014]; four participants in [van den Berg et al., 2016] and six participants in [Stine, Zylberberg, Ditterich, & Shadlen, 2020], assume that with extensive training, all participants would reach an acceptable level of performance. Accordingly, all participants received extensive training sessions on the Random-dot-motion discrimination task prior to data collection. To ensure that participants understood and followed the instruction correctly, they were asked to describe how they would complete the experiment and report their accuracy and confidence prior to experiments. In Experiment 1, one participant was excluded due to the difficulty in reporting decision and confidence simultaneously, and another participant withdrew from the experiment shortly after the start of the experiment. In addition, one participant was excluded from Experiment 2 because of the excessive noise in their EEG electrodes crucial to the analysis.

### 2.2 Stimuli

We explored confidence formation in a discrete environment with a random-dot-motion (RDM) discrimination task. Participants had to indicate the predominant motion direction of a cloud of moving dots (left or right) presented within a 5° circular aperture at the center of the screen. The dot density was 16.7 dots/degree^2^/s and the displacement of the coherently moving dots produced an apparent speed of 6 deg/s. The RDM movies were generated by three interleaved sets of dots presented on consecutive video frames. Three video frames later, each dot was redrawn at a location consistent with the direction of motion or a random location within the stimulus space. More details can be found in previous studies (e.g. [Roitman & Shadlen, 2002]). The experiment code was programmed in MATLAB 2016a (The Mathworks Inc., USA) using PsychToolbox (Brainard & Vision, 1997; Kleiner, Brainard, & Pelli, 2007).

### 2.3 Experimental Tasks

Participants performed the RDM task in blocks of 200 trials. Each trial started with participants fixating a small red point (diameter 0.3°) at the screen center. After 500 ms, two choice-targets appeared to the left and right of the fixation point (10° eccentricity; Figure 1A). Each target was shaped as a gradient rectangle (9° length and 0.5° width). After a variable duration of 200 - 500 ms (truncated exponential distribution), the RDM was presented. Participants had to indicate their choice after receiving one or two pulses of 120ms of motion pulses. The interpulse interval of double-pulse trials was selected randomly from 0, 120, 360, and 1080ms. On single-pulse trials, motion coherence was randomly selected from these six values: 0%, 3.2%, 6.4%, 12.8%, 25.6%, and 51.2%, whereas, on double-pulse trials, motion coherence of each pulse was randomly chosen from three values: 3.2%, 6.4%, and 12.8%. Both pulses had the same net direction of motion and participants were aware of it. In total, there were six single-pulse and 9 × 4 double-pulse trial types. After the offset of one or two motion pulse(s), a 400 to 1000 ms delay period (truncated exponential) was imposed before the Go signal appeared on the screen. In each trial, participants were required to indicate their response by directing the gaze to one of the targets, the upper extreme of targets representing full decision confidence and the lower extreme representing guessing (Figure 1A). We constructed a list of all possible conditions of motion coherences and gaps, then shuffled the listed conditions and assigned them randomly to the trials in each block to provide the approximate balance within the trials. Participants were instructed to achieve high performance. Distinctive auditory feedback (Beep Tones) was provided for correct and incorrect responses. The type of feedback of 0% coherence trials was selected randomly by a uniform distribution. In Experiment 1, each participant performed the task across multiple blocks on different days (12-20 blocks). Experiment 2 contained the same paradigm as Experiment 1. All variables of stimulus remained constant except, in Experiment 2, the EEG data were also recorded. In Experiment 2, each participant completed a session of 4-5 blocks.

### 2.4 EEG Recording and pre-processing

We used a 32-channel amplifier for the EEG signal recording (eWave, produced by ScienceBeam, http://www.sciencebeam.com/) which provided 1K sample/s of time resolution. EEG was recorded at 31 scalp sites (Fp1, Fp2, AF3, AF4, C3, C4, P3, P4, O1, O2, F7, F8, T7, T8, P7, P8, FPz, Fz, Cz, Pz, Oz, POz, FC1, FC2, CP1, CP2, FC5, FC6, CP5, CP6). The EEG signals were referenced to the right mastoid. The recorded data were taken to Matlab (The Mathworks Inc., USA) and pre-processed as follows. The signals were filtered using a band-pass filter from 0.1 Hz to 40 Hz (Zizlsperger et al., 2014) for removing high frequency and independent cognitive noises. Then, all trials were inspected, and those containing Electromyography (EMG) or other artifacts were identified and manually removed. The second artifact rejection step included independent components analysis (ICA) using the EEGLAB toolbox (Delorme & Makeig, 2004). To select the removable ICA component, the ADJUST plugin (Mognon, Jovicich, Bruzzone, & Buiatti, 2011) was used.

### 2.5 Pupillometry Recording and pre-processing

The eye data were collected using an EyeLink 1000 infrared eye-tracker system (SR Research Ltd. Ontario, Canada). This device allowed a 1000-Hz sampling rate and was controlled by a dedicated host PC. The system was calibrated and validated before each block by presenting nine targets at the display monitor’s center, edges, and corners of the. The left eye’s data was recorded and passed to the host PC via an Ethernet link during data collection.

Missing data and blinks, as detected by the EyeLink software, were padded and interpolated. Additional blinks were spotted using peak detection on the pupil signal’s velocity and then linearly interpolated (Mathôt, 2013).

### 2.6 Experimental procedure

Participants were given a consent form in which the experiment was described in general terms. After providing written informed consent, participants completed the tasks in a semidark, sound-attenuating room to minimize distraction in both experiments. All instructions were presented and stimuli were displayed on a CRT monitor (17 inches; PF790; refresh rate, 75 Hz; screen resolution, 800 × 600). A head and chin rest confirmed that the distance between the participants’ eyes and the monitor’s screen was 57 cm throughout the experiments. Participants were presented with demographic questions followed by training sessions and main sessions, respectively. The experimental protocol was approved by the ethics committee of the Iran University of Medical Sciences.

### 2.7 Data Analysis

Data analysis was performed using Matlab 2019a (The MathWorks Inc., United States).

#### 2.7.1 Behavioral analyses

Except where otherwise specified, we reported behavioral data of the first experiment; however, all the analyses were repeated for the EEG experiment and if the results were inconsistent, it has been admitted (Experiment 2 results were reported in Supplementary Figures 1, 2, 3, 4, 5, and 9).

We used linear and logistic regression models to evaluate the influence of the stimulus characteristics on choice, and confidence and to evaluate the null hypothesis that a single coefficient of a linear regression model is zero we ran *t*-tests. For logistic regression models, we used maximum likelihood under a binomial error model (i.e., a GLM) to evaluate the null hypothesis that one or more of the regression coefficients were equal to zero. *P_correct_* was the probability of correct response and *Logit*[*P_correct_*] indicated 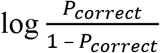. Also, *P_high_* was the probability of high confidence, *Logit*[*P_high_*] indicated 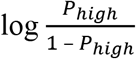 and *β_i_* denoted fitted coefficients

For single-pulse trials, to evaluate whether choice accuracy improved with coherence, we used the following logistic regression:

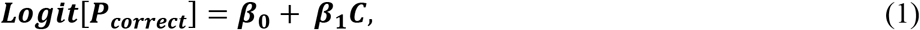

where *C* was motion strengths of the stimulus. The null hypotheses were that coherence would have no effect on choice accuracy (*H*_0_: *β*_1_ = 0). For single-pulse trials, we also used the following function to determine whether confidence improved consistently with coherence and accurate choices:

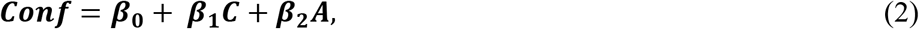

where *C* was motion strengths of the pulse and *A* was the accuracy of the response (0 or 1 for incorrect and correct). The null hypotheses were that coherence and accuracy would have no effect on confidence (*H*_0_: *β*_1–2_ = 0).

We also used linear regression to evaluate the effect of interpulse interval on confidence in double-pulse trials:

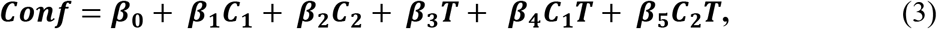

where *C*_1_ and *C*_2_ were motion strengths of each pulse, and *T* was the interpulse time interval. For double-pulse trials with equal pulse strength (*C*_1_ = *C*_2_), the redundant regression terms (*β*_2_, *β*_4_) were omitted. The null hypothesis was that the interpulse interval would not affect reported confidence (*H*_0_: *β*_3–5_ = 0). The similar equation was used to assess the relation of accuracy and the interpulse interval. To evaluate the impact of pulse sequence on confidence, the following regression model was fitted:

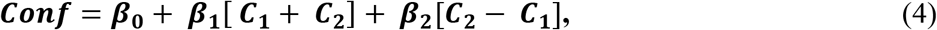

where *C*_1_ and *C*_2_ were corresponding motion strengths of each pulse. *β*_2_ indicated how the confidence varied from trials in which *C*_1_ > *C*_2_ to trials with a reversed sequence of motion pulses *C*_1_ < *C*_2_. The null hypothesis was that the sequence of motion pulses did not influence the confidence (*H*_0_: *β*_2_ = 0).

In addition, to investigate the variation of confidence in double-pulse trials compared to single-pulse trials, we subtracted confidence of double-pulse trials from corresponding confidence in single-pulse trials for each participant. For example, the confidence of a sequence of 3.2%, 6.4% motion strength trial, subtract separately once from 3.2% and once from 6.4% corresponding confidence in single-pulse trials. The process repeated for the data of correct and incorrect trials too. Moreover, the same method was used to compare the accuracy of double-pulse and single-pulse trials. To assess the effect of choice accuracy on the variation of confidence in double-pulse and single-pulse trials, we fitted the following:

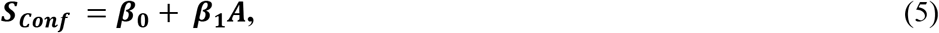

where the *S_Conf_* was the subtraction of confidence in double-pulse trials from corresponding single-pulse trials and *A* was the accuracy of the response (0 or 1 for incorrect and correct). The null hypothesis was the choice accuracy did not affect the variation of *S_Conf_* (*H*_0_: *β*_1_ = 0).

Moreover, we compared accuracy and confidence between single-pulse and double-pulse trials directly by fitted linear regression models using indicator variables for type of trials (1 for single-pulse and 0 for double-pulse). Also, as a sensitivity analysis we randomly chose the same numbers of trials from each coherence of single and double pulse trials of each participants’ data where the reported confidence was between 40 to 60. Then, we compared the probability of correct choice — accuracy—of the selected data. This range was chosen to make sure there were enough numbers of trials for all conditions.

##### 2.7.1.1 Response-time analysis

In the current study, response-time was referred to the time interval between the response cue onset and the participant’s response. To represent the trace of high and low confidence from behavioral data, we have selected an equal number of single/double-pulse trials from each participant’s data for 100 times. Then individual response-time were rank-ordered and binned into four quintiles. Next, we calculated he accuracy of high and low confidence trials in each bin. We expected to see a noteworthy difference between the accuracy of each bin grouped by levels of confidence. We only included motion strength of 3.2%, 6.4%, 12.8% of single-pulse trials (similar to coherence used in double-pulse trials) to control the impact of coherence on response-time. We used a linear model with separate intercepts for each participant on response-time to check whether the difference of response-time in high and low confidence can be traced in both single-pulse and double-pulse trials:

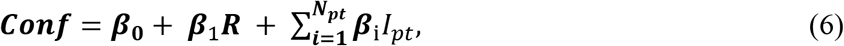

where *R* was the response-time of each trial and *I_pt_* is an indicator variable that takes a value of 1 if the trial was completed by participant *pt* and 0 otherwise. The null hypothesis was that confidence did not depend on the response-time (*H*_0_: *β*_1_ = 0). Moreover, as a sensitivity analysis to evaluate the relation of delay-time imposed before the cue onset and response-time, we used a linear regression model to see if the delay times would predict the response-time. The null hypothesis was that the response-time did not rest on the delay-time.

#### 2.7.2 Motion energy analysis

Random dot stimulus is stochastic, so the sensory evidence fluctuated within and across trials but around the nominal coherence level. To examine the fluctuations in motion during each trial, we filtered the sequence of random dots by using two pairs of quadrature spatiotemporal filters, as specified in previous studies (Adelson & Bergen, 1985; Kiani, Hanks, & Shadlen, 2008; Zylberberg et al., 2012). Since we aimed to understand the temporal course of confidence, we summed the energies across trials for each pulse in single/double-pulse trials.

We used a linear regression to test whether the confidence was more influenced by the second pulse’s motion energy than that of the first pulse in double-pulse trials. We tested double-pulse trials with unequal motion strength using the following linear regression model:

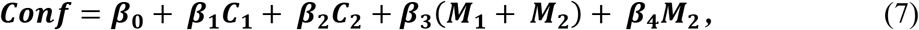

where *M*_1_ and *M*_2_ were the motion energy of each pulse and *C*_1_, *C*_2_ were corresponding motion strengths of each pulse. The null hypothesis was (*H*_0_: *β*_4_ = 0). For double-pulse trials with equal pulse strength (*C*_1_ = *C*_2_), the redundant regression terms (*β*_2_) were omitted. We used only double-pulse trials with nonzero interpulse interval in this analysis to avoid overlap in the windows for the first and second pulse. To certify that our findings were not relevant to participant performance, we analyzed motion energy in the two confidence levels only for correct responses. To evaluate the relation of confidence and motion energy in single-pulse trials, we fitted a linear regression model as follows:

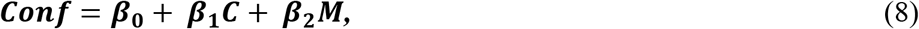

where *M* was the motion energy of the presented motion stimulus, *C* was the pulse coherence and the null hypothesis was that confidence did not depend on the motion energy (*H*_0_: *β*_2_ = 0).

#### 2.7.3 General computational modeling approach

We implemented a set of computational models based on signal detection theory to provide a mechanistic explanation of the experimental data. According to SDT, observers set a decision criterion (*cr*) to discriminate between two stimuli (e.g., labeled as *S*_1_ and *S*_2_). They also set criteria *cr*_2,“*S*1”_ and *cr*_2,“*S*2”_ to determine confidence ratings around the decision criterion *cr* (Figure 1B; for more details, see Supplementary Appendix 2). In order to fit confidence data to SDT, we categorized the confidence ratings as either high or low. Since the participants were told to choose the upper part of the bar as high confidence and the lower part as low confidence (ranged from 0 to 100), we considered reported confidence higher than midline as high confidence and lower than midline as low confidence respectively. However, in addition to the midline, we tested various binary level set methods for categorizing participants’ high and low confidence ratings. First, the highest 55% and 45% of each participant’s confidence reports were considered high confidence (similar to [Zylberberg, Wolpert, & Shadlen, 2018]). Then, the mean of each participant’s confidence was calculated separately, and the confidence ratings above the mean were considered high ratings. Using these methods did not significantly alter reported confidence categorization (see Supplementary Figure 8).

We computed stimulus sensitivity (*d*’), confidence criteria, and the measures of metacognitive ability (*Meta*–*d*′, *Meta*–*d*’/*d*’) of single-pulse and double-pulse trials. We used code provided by Maniscalco and Lau (Maniscalco & Lau, 2012) in which metacognitive sensitivity (*Meta*–*d*′) is computed by setting the *d*′ value that would produce the observed confidence. In addition, *Meta*–*d*’/*d*’ was calculated by normalizing *Meta*–*d*’ by *d*′ through division. To support the fact that our findings were not relevant to variation of coherence of single and double-pulse trials, we only included single-pulse trials with motion strength of 3.2%, 6.4%, 12.8%. In addition, we fitted the SDT model with trials simulated by a perfect integrator model (the model is described later). However, one difference between groups was that they might not be matched for the number of trials: the single-pulse included on average fewer trials for each coherence per participant compared to double-pulse trials. Previous research has suggested that the number of trials could bias measures of metacognitive ability (Fleming, 2017). Therefore, in a control analysis, we created 100 sets of trials randomly from the single/double-pulse trials and from trials simulated by the perfect integrator model. Each set contained the same number of trials for each participant. We then averaged the metacognitive scores obtained from these 100 sets and repeated the comparison procedure.

#### 2.7.4 Perfect integrator Model

To estimate the expected confidence (*P*_*e*(*high*)_) in double-pulses trials, we assumed that each trial’s confidence was achieved based on evidence integrating from both pulses by using a perfect integrator model. In the perfect integrator model, the expected accuracy (*P*_*e*(*correct*)_) for double-pulse trials computed as the following (Kiani et al., 2013):

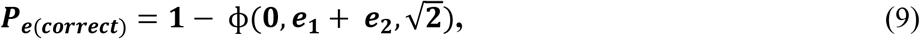

where *e*_1_ and *e*_2_ were the pieces of evidence that underlie by *P*_1_ and *P*_2_ (the probabilities of the correct choices in corresponding single-pulse trials) and were computed as:

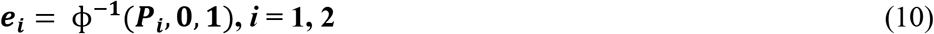

where ϕ^−1^ was inverse ϕ, which represented the cumulative Gaussian distribution (Kiani et al., 2013).

To predict the confidence of double-pulse trials by this model, *cr* and *d*′ were computed based on integrated evidence (Eq.9, 10, for more details see Supplementary Appendix 2). Then, confidence on correct and error trials were calculated based on confidence performance from corresponding single-pulse trials (similar to Eq.9, 10). Accordingly, confidence (for both correct response or incorrect response) would be predicted by the perfect integrator model. Besides, the model parameters, including confidence criteria along with *d*′, *Meta*–*d*′ and, *Meta*–*d*′/*d*′ were computed.

#### 2.7.5 Optimized models

We considered two optimized models. The purpose of these two models was to address how improving stimulus or metacognitive sensitivity could influence confidence prediction. In the type 1 optimized model, we optimized the perfect integrator model by letting the difference of *d*’ (the measures of task performance in SDT) from the perfect integrator and double-pulse trials minimized for each participant using the fmincon function in the optimization toolbox for Matlab. In the type 2 optimized model, using the similar optimization technique, we optimized the *Meta*–*d*′ in the perfect integrator model by providing each participant’s *Meta*–*d*′ computed from behavioral data. In other words, in type 1 optimized model, stimulus sensitivity was optimized, whereas in the type 2 optimized model, metacognitive sensitivity was optimized. Thus, we examined how improving stimulus or metacognitive sensitivity could influence confidence prediction.

#### 2.7.6 Model evaluation

We evaluated the models qualitatively (i.e., parameter recovery exercises) and quantitatively (i.e., maximum likelihood estimation).

In the qualitative method, based on the calculated parameters of the model, the probability of choosing high confidence for all combinations of motion strength for each participant was calculated ([Maniscalco & Lau, 2014]; see Supplementary Appendix 2). We compared the expected high confidence predicted by models to the observed high confidence in double-pulse trials using a linear regression model, as follows:

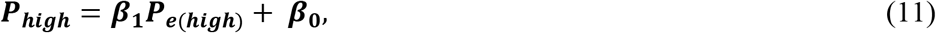

where *P*_*e*(*high*)_ was the expected probability of high confidence while ***P_high_*** was observed high confidence in double-pulse trials. We compared slope (*β*_1_) against the 1:1 line in each model.

In addition, to compare models quantitatively, an equal number of trials from each participant’s trials was selected randomly and then each model fitted to the selected data. This procedure was repeated for 100 times, then the computed MLEs of each model were averaged.

#### 2.7.7 Confidence suboptimality

The optimal decision-making is disrupted by several sources of suboptimality (Balsdon et al., 2020). In SDT, an added noise, *ξ_n_*, represents a potential loss of information between sensory decision information and metacognitive information, such as confidence rating. This noise has a Gaussian distribution with zero mean and standard deviation *σ* (Maniscalco & Lau, 2014). The parameter *σ* determines how much the metacognitive variable is noisier than the decision variable (Maniscalco & Lau, 2014).

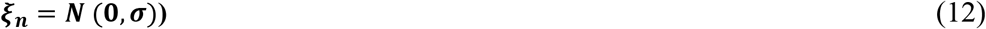

This noise is related to metacognitive efficiency —*Meta*–*d*′/*d*′ (Maniscalco & Lau, 2014). We simulated trials using the same parameter values resulting from the perfect integrator model, except for increased noise, to consider this confidence suboptimality.

#### 2.7.8 EEG analysis

In current study, the EEG analysis focused on a neural marker of perceptual decision-making linked with stimulus preparation and stimulus processing. First, we epoched the EEG responses for averages over the entire trial for all coherence. To characterize the time-course of confidence in the ERP data, time courses were baseline-corrected throughout 100 ms before stimulus onset and split by high and low confidence. Next, the component we focused on was the centro-parietal positivity (CPP) which possibly is identical to the classic P300 component (Herding et al., 2019; Twomey, Murphy, Kelly, & O’connell, 2015). The CPP is associated with the sampling of available evidence in perceptual decisions and confidence rating at the time period of 200-500 ms after stimulus onset (Herding et al., 2019; Rausch, Zehetleitner, Steinhauser, & Maier, 2020; Vafaei Shooshtari et al., 2019; Zizlsperger et al., 2014) or at the time of the response (Boldt et al., 2019). In current study, CPP amplitude was measured as the mean amplitude in a time-window ranging from 200 ms to 500 ms after stimulus onset in an electrode cluster containing the electrodes CP1, CP2, Cz, and Pz (Boldt et al., 2019; Herding et al., 2019; Rausch et al., 2020; Tagliabue et al., 2019; Twomey, Kelly, & O’Connell, 2016; Vafaei Shooshtari et al., 2019; Zizlsperger et al., 2014). We epoched the EEG responses aligned to the stimulus onset, from 200 ms pre-stimulus to 500 ms post-stimulus for each pulse. Then, these epochs were baselined to a window −100 ms to stimulus-locked to prevent differences in the visual response to the stimulus affecting the baseline. To avoid overlap in the windows for the first and second pulse, we used only double-pulse trials with 360 and 1080 ms interpulse interval in this analysis.

The ERP signals were examined for levels of confidence separately in double-pulse trials and single-pulse trials. To support the fact that our findings were not relevant to motion pulse strength in double-pulse trials, we tested double-pulse trials with equal motion strength pulses. To test if CPP amplitude could distinguish between high and low confidence trials, we performed an independent samples *t*-test comparing the average of the high and low confidence trials at each time sample on the EEG signal of the four CPP electrodes at a significance threshold corresponding to a p-value of 0.05. Correction for multiple comparisons was performed using a cluster-based method (Maris and Oostenveld, 2007). Each cluster was characterized by the sum of the t-statistics comparing the average high and low confidence CPPs. These summed t-statistics were then compared to clusters of t-statistics obtained from a Monte Carlo simulation with 3000 iterations. To make sure that our findings were not relevant to participants’ performance, as a secondary check, we analyzed the CPP signals in two confidence levels only for correct trials.

To present a statistical comparison of the single-pulse and double-pulse trials and to examine whether CPP amplitude could predict confidence for both single-pulse and double-pulse trials, we averaged the time courses of electrodes CP1, CP2, Cz, and Pz and then performed a linear regression between the amplitude of this signal of each participant and their mean confidence:

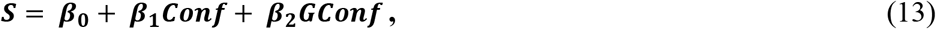

where *S* was the mean amplitude of time period of 200-500 ms after stimulus onset for each participant and *G* was an indicator variable that took a value of 1 if the trial was double- and 0 otherwise. The null hypothesis was that the type of trials did not affect the impact between confidence and CPP signal (*H*_0_: *β*_2_ = 0).

#### 2.7.9 Pupillometry analysis

First, to characterize the time-course of confidence in the pupillometry data, time courses were baseline-corrected throughout 200 ms before stimulus onset and split by high and low confidence. Previous studies showed that pupil dilation response after choice and before feedback reflected decision uncertainty (Colizoli, De Gee, Urai, & Donner, 2018; Urai et al., 2017). In current study, as confidence is an uncertainty complement (Hebart, Schriever, Donner, & Haynes, 2014; Kepecs & Mainen, 2012), the method was implemented to study the relation of confidence and pupil dilation response. Accordingly, the baseline-corrected pupil signal throughout 200 ms before feedback was calculated as our single-trial measure of pupil response. We epoched trials and baselined each trial by subtracting the mean pupil diameter 50 ms before the response. We included all trials of both experiments in the analyses reported in this paper. To control the impact of coherence on pupil response, we only included motion strength of 3.2%, 6.4%, 12.8% of single-pulse trials.

According to the temporal low-pass characteristics of the slow peripheral pupil apparatus (Hoeks & Ellenbroek, 1993) even in the absence of amplitude variations in the underlying neural responses, trial-to-trial variations in response-time can impact pupil responses (Urai et al., 2017). Therefore, we removed the trial-to-trial fluctuations in pupil responses via a linear regression (De Gee, Urai, & Donner, 2018; Urai et al., 2017) and used it for analyses reported in this study. Statistical inferences were performed using a cluster-based procedure to examine whether pupil response could distinguish between high and low confidence trials (Maris and Oostenveld, 2007). Significant clusters were found using a *t*-test at each time-point (at a statistical threshold of p < 0.05) and comparing the sum of the t-statistics in these clusters to those obtained over 3000 permutations. As a secondary check, we analyzed the pupil response signals in two confidence levels only for correct trials to support the fact that our findings were not relevant to participants’ performance.

To present a statistical comparison of the single-pulse and double-pulse trials and to examine whether pupil response could predict confidence for both single-pulse and double-pulse trials, we performed a linear regression between the average of pupil responses for each participant and their mean confidence:

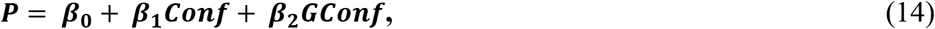

where *P* was mean the average of pupil response for each participant and *G* was an indicator variable that took a value of 1 if the trial is double-pulse trials and 0 otherwise. The null hypothesis was that the type of trials did not affect the relationship between confidence and pupil response signal (*H*_0_: *β*_2_ = 0).

#### 2.7.10 General statistical analysis

To test our hypotheses, a series of regression analyses were run after confirming the assumptions of linear and logistic regression were met. Effect sizes were reported and as suggested, here, we considered small (*f^2^* = .02), medium (*f^2^* = .15), and large (*f^2^* = .35) effect sizes (see [Cohen, 1970]) at the alpha level of 5%.

Moreover, we used repeated-measures *t*-tests. As suggested, we considered small (d = .2), medium (d = .5), and large (d = .8) effect sizes for this assessment (see [Cohen, 1970]) and the statistical significance for *t*-tests was set to 0.05.

Throughout the analysis, Bonferroni corrections were applied to p-values less than 0.05 when more than one statistical test was carried out per hypothesis (with this indicated by bonf*[number of tests corrected for] in subscript), whereas non-significant p-values were reported uncorrected.

For tests of pupil response signals and ERPs between two levels of confidence, statistical inferences were performed using a cluster-based procedure (Maris, E. & Oostenveld, R, 2007). All cluster-based permutations were done using the FieldTrip toolbox (Oostenveld et al., 2011) in MATLAB (at a statistical threshold of p < 0.05).

## 3 Results

We tested our predictions in two studies that applied the same paradigm (Figure 1A). The first study used behavioral measures and pupillometry analyses, whereas for the second experiment, we recorded EEG signals as well. Participants decided about the direction of the RDM motion based on brief motion pulses. The task design contained different conditions which allowed us to compare participants’ behavior in (i) one or two brief pulse(s) of random dot motion, (ii) different coherence of motion stimulus, and (iii) four distinct interpulse intervals.

### 3.1 Behavioral results

We used the single-pulse trials to benchmark the effect of coherence on choice accuracy and confidence. As shown in Figure 2A, for single-pulse trials, participants were more confident for high coherence stimuli (Figure 2A; Eq.2; *β_1_* = .06, *p_bonf*2_* < .001, 95% CI = [.04, .08], *f^2^* = .23), ranged from 43 for 3.2% to 96 for 51.2% coherence. Also, accuracy improved with motion strength reached from 56% for 3.2% to 99% for 51.2% (Figure 2B, black line; Eq.1; *β_1_* = .10, *p* < .001, 95% CI = [.08, .12], *f^2^* = .36). In general, they also had better performance whenever they had reported higher confidence comparing to lower confidence (Figure 2B, red and green; Eq.2; *β_2_* = .12, *p_bonf*2_* < .001, 95% CI = [.08, 0.15], *f^2^* = .23) except in weak coherence (3.2%) that the difference of high and low confidence was not significant (Figure 2B, red and green; Eq.2; *β*_2_ =.01, *p* = .85, 95% CI = [−.07, 0.08], *f^2^* = .01).

**Figure 2.**
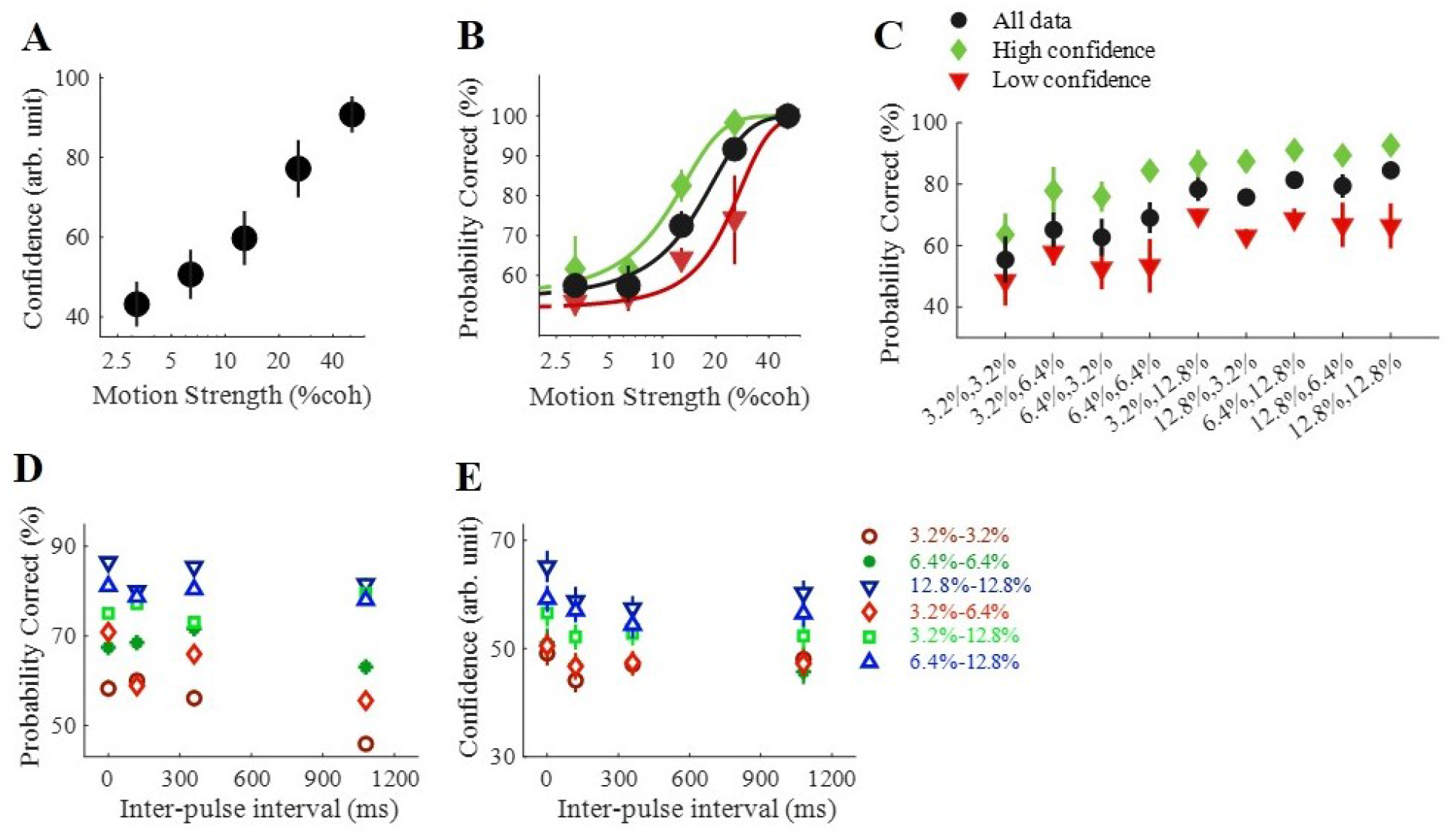
The interplay between confidence, accuracy, and coherence in single/double-pulse trials, and the interpulse interval in double-pulse trials. **(A)** Average confidence reported as a function of the motion coherence in single-pulse trials. **(B)** Accuracy in single-pulse trials and **(C)** double-pulse trials in all trials (black), split by high (green) and low (red) confidence decisions. **(D)** Choice accuracy for double-pulse trials grouping in all possible interpulse intervals. **(E)** Confidence of double-pulse trials was calculated by pooling data across all interpulse intervals. In (B) curves are model fits. In (D) and (E) each data point reports pooled data from indicated sequence pulse and its reverse order (e.g., 12.8–3.2% and 3.2 −12.8%). In all panels, data are represented as group mean ± SEM.

Moreover, in double-pulse trials, the accuracy improved with motion strength (Figure 2C, black dots) and participants were more accurate while reporting higher confidence (Figure 2C, green dots). Along with accuracy (Figure 2D; all *ps* > .1, [Kiani et al., 2013; Tohidi-Moghaddam et al., 2019], see Supplementary Figure 1A for Experiment 2 data), the confidence was largely unaffected by interpulse interval in double-pulse trials (Figure 2E; Eq.3; all *ps* > .1; Supplementary Table 2, results of individual participants; see Supplementary Figure 1B for Experiment 2 data). The two pulses separated by up to 1 s supported a level of confidence that was indistinguishable from a pair of pulses separated by no gap. To make sure that the trials with zero interpulse interval did not differ from trials with nonzero interpulse intervals, similar analyses were performed on data including trials with zero interpulse interval and just one of the nonzero interpulse intervals. Results revealed no significant difference (Eq.3, all *ps* > .1; Supplementary Table 3). Further, confidence was unaffected by the interpulse interval for double-pulse trials across each possible coherence sequences (Supplementary Table 4).

Direct comparison between participants’ accuracy of single-pulse and double-pulse trials, along with previous studies (Kiani et al., 2013; Tohidi-Moghaddam et al., 2019), showed that participants’ accuracy significantly differed (*β* = .09, *p*< .001, 95% CI = [.08, .11], *f^2^* = .21). However, in double-pulse trials participants were not more confident compared to single-pulse trials (*β* = .02, *p* = .15, 95% CI = [−.01, .04], *f^2^* = .01).

Furthermore, although the order of the pulses affected accuracy (Figure 3A, [Kiani et al., 2013; Tohidi-Moghaddam et al., 2019]), participants were not more confident in double-pulse trials with unequal pulse strength where the stronger motion appeared in a second order (Figure 3B; Eq. 4; *β_2_* = .01, *p* = .51, 95% CI = [.00, .02], *f^2^* = .01; see Supplementary Figure 2B for Experiment 2 data). Also, we explored whether if the first pulse would have less of an impact on confidence after a longer delay. According to the results, the order of the pulses failed to show significant effects on confidence on any of interpulse intervals (Supplementary Table 5).

**Figure 3.**
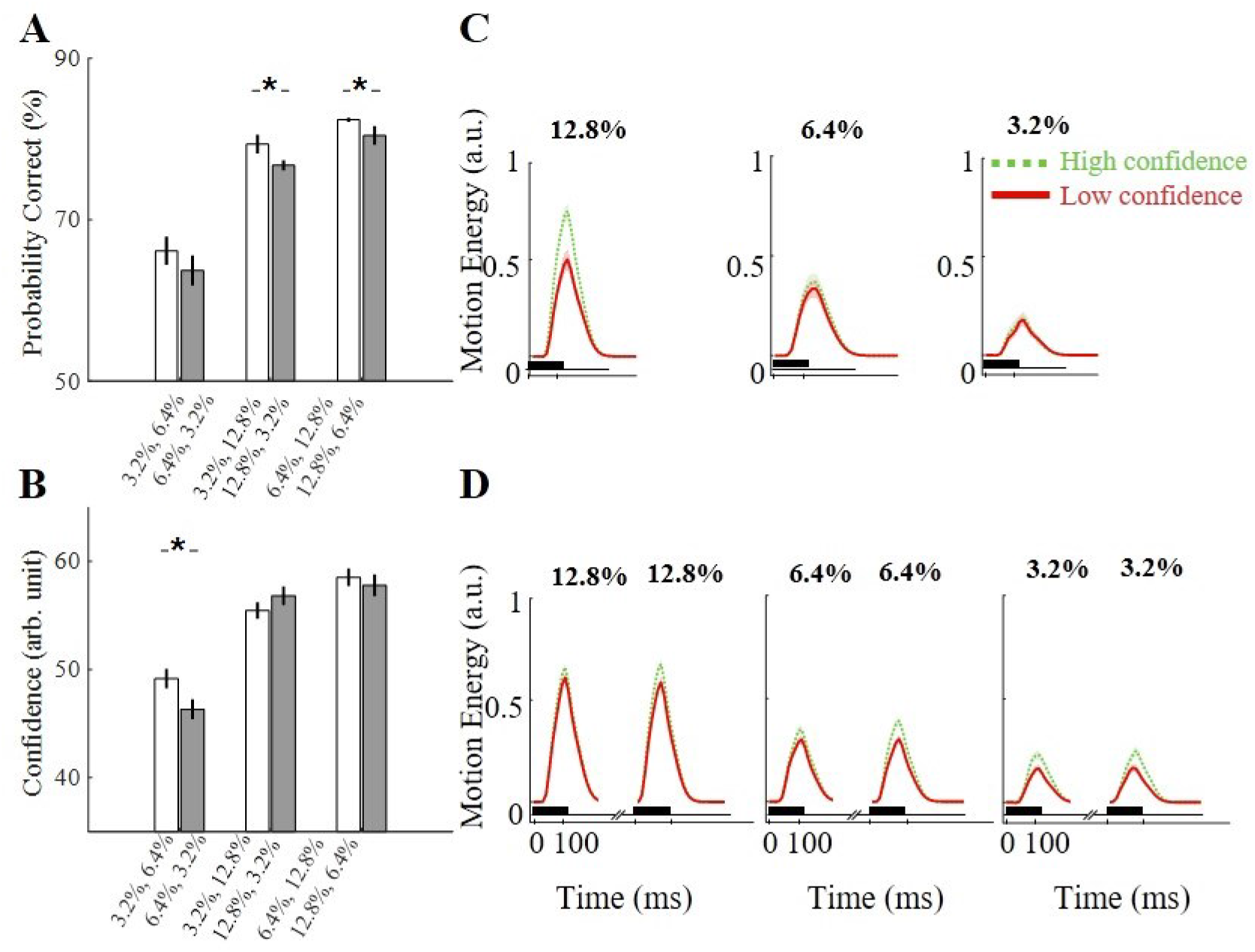
Choice confidence did not depend on the sequence of motion pulses. **(A)** The weak–strong pulse sequence contributed higher accuracy than the strong–weak sequence. Accuracy for double-pulse trials with unequal pulse strength was calculated by pooling data across all intertrial intervals. **(B)** The weak–strong pulse sequence did not contribute higher confidence than the strong–weak sequence. In all panels, data are represented as group mean ± SEM. (*p<0.05) **(C)** In single-pulse trials, low and high confidence cannot be determined by motion energy profiles in weaker pulses **(D)** The second pulse had slightly more impact on confidence. Data were pooled for all nonzero interpulse intervals while only correct trials with equal pulse strength are included. In (C) and (D), the shaded region around the mean indicates SEM. The black horizontal bars show the duration of the stimulus display. The units of motion energy are arbitrary and the same for all motion strengths.

### 3.2 Motion energy results

To yield a precise estimate of the decision-relevant sensory evidence accommodated in the stochastic stimuli, we employed motion energy filtering to the random dot motion stimuli. Figure 3D displays the average motion energy in double-pulse trials with equal pulse strength. A linear regression confirmed the influence of trial-to-trial fluctuations of motion energy on confidence (Eq.7; *β_2_* = .10, *p_bonf*4_* < .001, 95% CI = [.04, .16], *f^2^* = .19). However, the difference of the motion energy profiles for high and low confidence responses was not larger for the second pulse than the first pulse with equal pulse strength (Eq.7; *β_3_* = .11, *p_bonf*4_* = .15, 95% CI = [.07, .15], *f^2^* = .13), as well as, with unequal pulse strength (Eq.7; *β_4_* = .10, *p* = .06, 95% CI = [.06, .14], *f^2^* = .08). Consequently, motion energy analysis could not provide independent confirmation of the asymmetric effect of pulses for confidence.

Additionally, in single-pulse trials, the motion energy profiles for high and low confidence with stronger pulse strength (12.8%, 6.4%) were significantly different (Figure 3C; Eq.8; *β_2_* = .41, *p_bonf*2_* < .001, 95% CI = [.23, .59], *f^2^* = .24). However, the motion energy profiles in weak motion pulses were not significantly different (Figure 3D; Eq.8; β2 = .17, p = .44, 95% CI = [−.26, .60], f2 = .10). Accordingly, motion energy analysis suggests that when the pulses’ motion strengths are weak, the participants’ confidence would not depend on the motion energy.

### 3.3 The Interplay between confidence in single vs. double pulse(s) trials

To address accuracy and confidence variation in double-pulse from single-pulse trials, we subtracted *P_correct_* or confidence of each coherence (3.2%, 6.4% and 12.8%) in single-pulse trials from *P_correct_* or confidence of corresponding sequence in double-pulse trials. As we expected, *P_correct_* improved in all coherence combinations (Figure 4A). Additionally, considering all the trials, confidence increased when the other pulse was a strong pulse (12.8%) (Figure 4B and Figure 4C for correct trials), while decreased or did not change whenever the other pulse was a weak motion strength (3.2%, 6.4%). Interestingly, in error trials, confidence on double-pulse trials decreased compared to single-pulse trials for all conditions (Figure 4D). These results did not influence by the duration of interpulse intervals (Supplementary Figure 6A, B, C, D).

**Figure 4.**
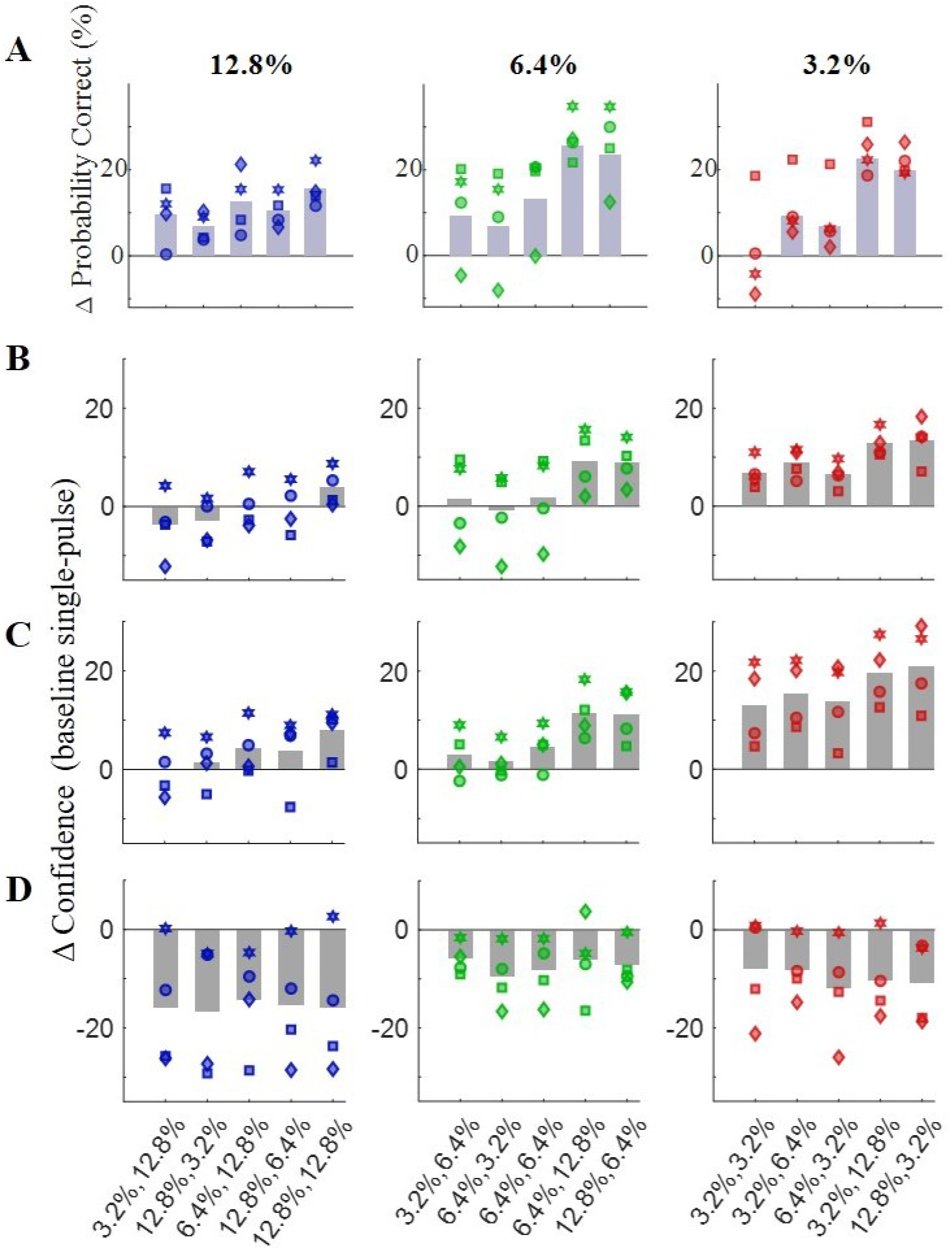
The variation of accuracy or confidence in double-pulse trials baselined by single-pulse trials. **(A)** Considering all the trials, the accuracy improved in most of the combinations in double-pulse trials. **(B)** Considering all the trials, the confidence improved in the combination with stronger pulses while the confidence in sequence with a weaker pulse either decreased or remained constant. **(C)** In correct-choice trials, the increasing effect of stronger pulses was enhanced and the confidence even slightly improved in combination with weaker pulses compared to corresponding baseline. **(D)** Interestingly, in incorrect trials, the confidence decreased in every condition. The gray bars represent mean data while each dot shows the data of each participant.

In other words, the participants reported lower confidence in double-pulse trials compared to single-pulse trials for incorrect choices but reported higher confidence for correct choices (Figure 4 Eq.5, *β_1_* = .15, *p* < .001, 95% CI = [.13, .17], *f^2^* = .29; Supplementary Figure 3 for Experiment 2; Supplementary Table 1, results of individual participants). This data is in line with the fact that good metacognitive sensitivity will provide higher confidence for correct responses, and lower for incorrect ones (Fleming, 2017; Maniscalco & Lau, 2012, 2014). Accordingly, we implemented computational models in order to study the relation of accuracy and confidence (confidence resolution) in single-pulse and double-pulse trials.

### 3.4 Computational models

In SDT models, *d*′ is stimulus sensitivity and has relation to the choice performance. In general, as we expected, *d*′ in double-pulse trials surpasses the *d*′ in single-pulse trials (Figure 5C left, Supplementary Figure 4 left). Furthermore, compared to single-pulse trials, metacognitive sensitivity, *Meta*–*d*′, increased in double-pulse trials (Figure 5C middle, Supplementary Figure 4 middle). We also computed metacognitive efficiency (*Meta*–*d*′/*d*′), as another index of the metacognitive ability which attempt to equate for underlying accuracy. Here, *Meta*–*d*′/*d*′ computed from double-pulse trials was higher for most of the participants (Figure 5C right, Supplementary Figure 4 right). Pairwise comparisons of *d*′, *Meta*–*d*′, and *Meta*–*d*′/*d*′ across models was shown in Supplementary Table 6. Furthermore, *d*′, *Meta*–*d*′, and *Meta*–*d*′/*d*′ estimated for double-pulse trials were independent of the interpulse interval in both experiments (see Supplementary Figure 9 and Supplementary Table 7).

**Figure 5.**
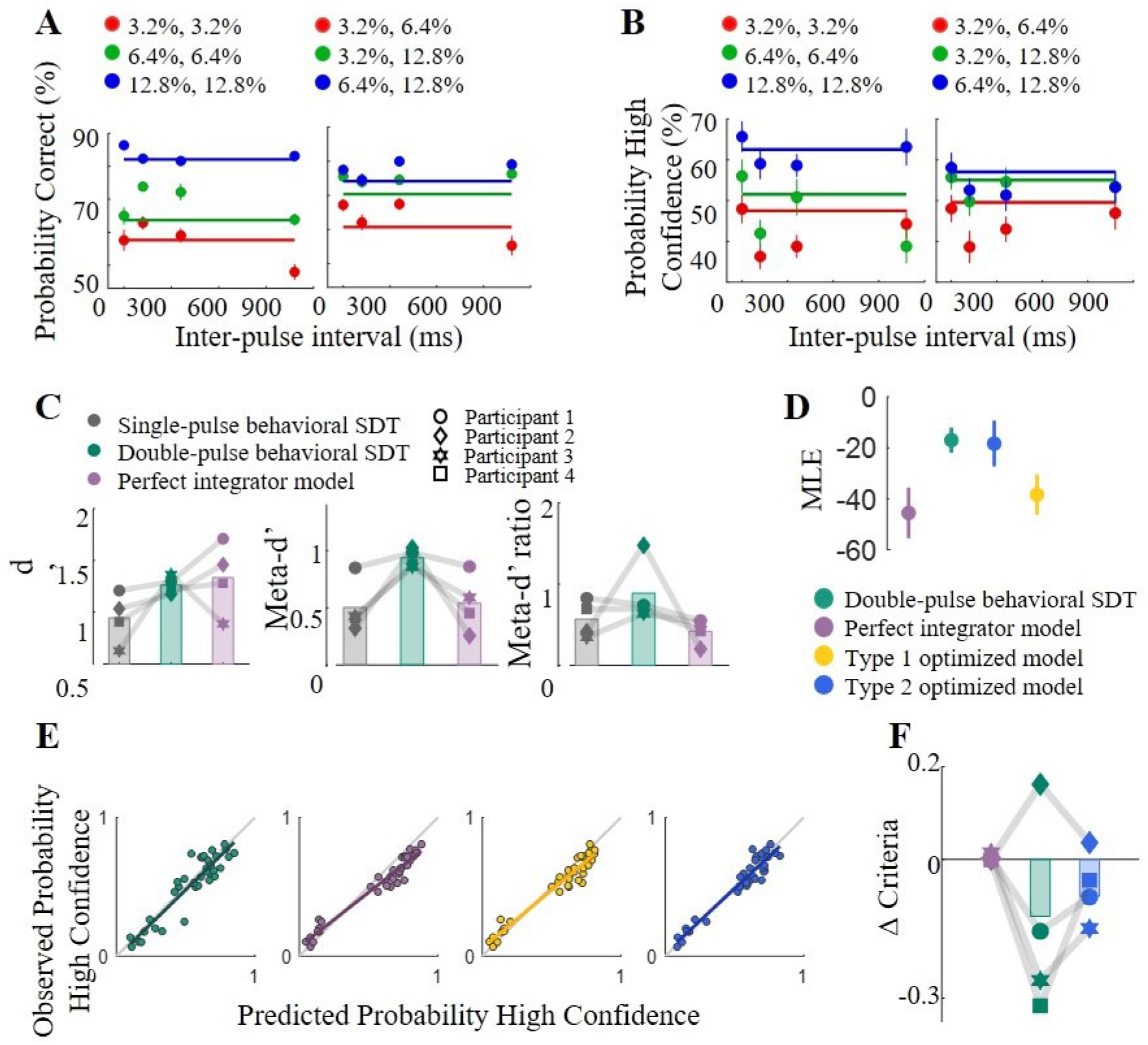
Comparison of the models and human behavior. **(A)** Accuracy in double-pulse trials. Horizontal lines show accuracy prediction by the perfect integrator model. **(B)** Confidence in double-pulse trials. Horizontal lines show confidence prediction by the perfect integrator model. In (A) and (B) each data point represents pooled data from the pulse sequence indicated by the legend and its reverse order. **(C)** Stimulus sensitivity (*d*′), metacognitive sensitivity (*Meta*–*d*′), metacognitive efficiency (*Meta*–*d*′/*d*′) estimated for single-pulse trials, double-pulse trials and the perfect integrator models for each participant. **(D)** Model comparison suggests strong evidence in favor of the type 2 optimized model over the perfect integrator model and the type 1 optimized model. **(E)** Relation of the observed confidence and the predicted confidence by simulating the model of double-pulse trials (green), the perfect integrator model (purple), the type 1 optimized model (yellow), and the type 2 optimized model (blue). Colored lines indicate the best-fitting slope of a linear regression analysis. Each data point represents pooled data from each sequence of pulses for each participant. **(F)** Variation of confidence criteria comparing to single-pulse trials in the perfect integrator model, the type 2 optimized model and, the model fitted to double-pulse trials. For panels A, B, and D data are represented as group mean ± SEM. For panels C and F, each dot represents the data of each participant

To further quantify the improvement of metacognitive ability in double-pulse trials —a more accurate appraisal of decisions—, we randomly chose the same numbers of trials from each coherence of single-pulse and double-pulse trials from each participants’ data where the reported confidence was between 40 to 60. Comparison among the probability of correct choice of these single-pulse and double-pulse trials showed higher performance in double-pulse trials with similar confidence ratings (see Supplementary Figure 7).

We estimated the expected accuracy and confidence that would be achieved in double-pulse trials under the assumption of perfect integration. The accuracy in double-pulse trials surpasses the expectation measured by the perfect integrator model (Figure 5A; [Kiani et al., 2013]). Considering the strong positive relation of accuracy and confidence (Kiani et al., 2014; Vafaei Shooshtari et al., 2019), we expected the observed confidence would exceed the predicted confidence calculated by the perfect integrator model (Eq.9), but it did not (Figure 5B). Further, we predicted the *d*′ in double-pulse trials by the perfect integrator model based on single-pulse trials performance. Accordingly, in general, *d*′ in double-pulse trials surpasses the predictions but if participants’ performance (*d*′) in single-pulse trials failed, the predicted performance for double-pulse trials missed the observed data (Figure 5C left, Supplementary Figure 4 left). Furthermore, in line with confidence, the perfect integrator model failed to imitate the *Meta*–*d*′ in double-pulse trials (Figure 5C middle, Supplementary Figure 4 middle). Likewise, *Meta*–*d*′/*d*′ simulated from the perfect integrator model failed to track *Meta*–*d*′/*d*′ observed in double-pulse trials (Figure 5C right, Supplementary Figure 4). Altogether, the perfect integrator failed to employ observed confidence and metacognitive ability in double-pulse trials.

As a control investigation, we examined whether the differences in estimated metacognitive ability between models could result from the different number of trials. We averaged the metacognitive scores obtained from randomly selected equal numbers of samples, and found very similar results. Thus, the difference in the estimated metacognitive efficiency cannot be explained by the difference in the number of trials between the single-pulse, double-pulse, and perfect integrator models (Supplementary Figure 10).

To sum, the perfect integrator model failed to follow observed metacognitive ability in double-pulse trials, and the *β*_1_ (slope in Eq.11) differed from 1:1 line which means predicted confidence failed to account for observed confidence (Figure 5E purple; Eq.11, *β_1_* = .77, *p* < .001, 95% CI = [.71, .83], *f^2^* = 15.66). Consequently, we introduced two models that optimized stimulus sensitivity (*d*′) or metacognitive sensitivity (*Meta–d′*) of the perfect integrator model. In type 1 optimized model, where the stimulus sensitivity (related to decision performance) optimized, the predicted confidence slightly improved (Figure 5E yellow, Eq.11, *β_1_* = .82, *p* < .001, 95% CI = [.79, .85], *f^2^* = 12.01). In the type 2 optimized model, the stimulus parameter (*d*′) remained similar to the perfect integrator model whereas the placement of confidence criteria was optimized to follow the *Meta*–*d*′ of the observed data. In this way, the predicted confidence improved considerably (Figure 5E blue, Eq.11, *β_1_* = .98, *p* < .001, 95% CI = [.88, 1.08], *f^2^* = 10.11). As an extra measure, we take the confidence criteria of the single-pulse SDT model as the baseline and measure the variation of criteria of the perfect integrator model, the behavioral model and the type 2 optimized model of double-pulse trials. The type 2 optimized model’s variation has changed compared to the perfect integrator model to follow the behavioral model (Figure 5F, and Supplementary Figure 5 for Experiment 2).

Accordingly, these results suggest the inability to predict the proper change in confidence criteria as a factor that made the perfect integrator model unable to estimate the confidence in double-pulse trials from single-pulse trials.

In addition, to investigate whether the suboptimality in confidence reporting and inability to predict the proper change in confidence criteria is related to confidence noise, we simulated data using the perfect integrator model’s parameters while setting higher confidence noise (Eq. 12). The predicted confidence from this simulation improved (Eq.11, *β_1_* = .97, *p* < .001, 95% CI = [.83, 1.07], *f^2^* = 9.00). Consequently, one problem of the perfect integrator model is ignoring the effect of confidence noise.

#### 3.4.1 Models’ evaluation

We conducted parameter recovery simulations to evaluate models. In single-pulse trials, linear regression indicated that there was a significant effect between the predicted and observed confidence, (Eq.11, *β_1_* = 1.04, *p* < .001, 95% CI = [.90, 1.18], *f^2^* = 8.09). In double-pulse trials, regression coefficient was statistically significant and close to 1:1 line (Figure 5E green; Eq. 11, *β_1_* = 1.03, *p* < .001, 95% CI = [.91, 1.15], *f^2^* = 11.05) meaning predicted confidence by classic SDT also explained a significant proportion of variance in the observed confidence. Moreover, as discussed before the type 2 optimized model intimate the behavioral data (Figure 5E blue), unlike the perfect integrator model (Figure 5E purple) and the type 1 optimized model (Figure 5E yellow).

A quantitative model comparison unsurprisingly favored the type 2 optimized model (mean MLE = −18.31) and the behavioral double-pulse model (mean MLE = −16.98) over the perfect integrator model (mean MLE = −45.53) (Figure 5D) and type 1 optimized model (mean MLE = −38.31).

### 3.5 Response-time analysis

As confidence was traced by both evidence and response-time in perceptual decision-making (Kiani et al., 2014; van den Berg et al., 2016; Zylberberg et al., 2016), we compared accuracy grouped by high and low confidence in four response-time bins. Accordingly, in double-pulse trials, the difference of accuracy grouped by high and low confidence was noteworthy in all response-time bins in double-pulse trials (Figure 6A) but not in single-pulse trials (Figure 6B). A linear regression confirmed the influence of response-time on confidence in double-pulse (*β_1_* = −.04, *p* < .01, 95% CI = [−.05, −.03], *f^2^* = .10) but not in single-pulse trials (*β_1_* = −.12, *p* = .62, 95% CI = [−1.78, −.07], *f^2^* = .04). Additionally, in our double-pulse trials, participants decided faster than single-pulse trials in all interval durations (Figure 6C). As a check, we regressed the delay-time before cue onset (0.4 to 1 s truncated exponential) and response-time in both single-pulse and double-pulse trials to examine the effect of imposed delay-time on response-time. The effect was small in both single-pulse (*β* = −.01 × 10^−5^, *p* = .004, 95% CI = [0.00, 0.00], *f^2^* = .00) and double-pulse trials (*β* = −.0003 × 10^−6^, *p* < .001, 95% CI = [0.00, 0.00], *f^2^* = .01).

**Figure 6.**
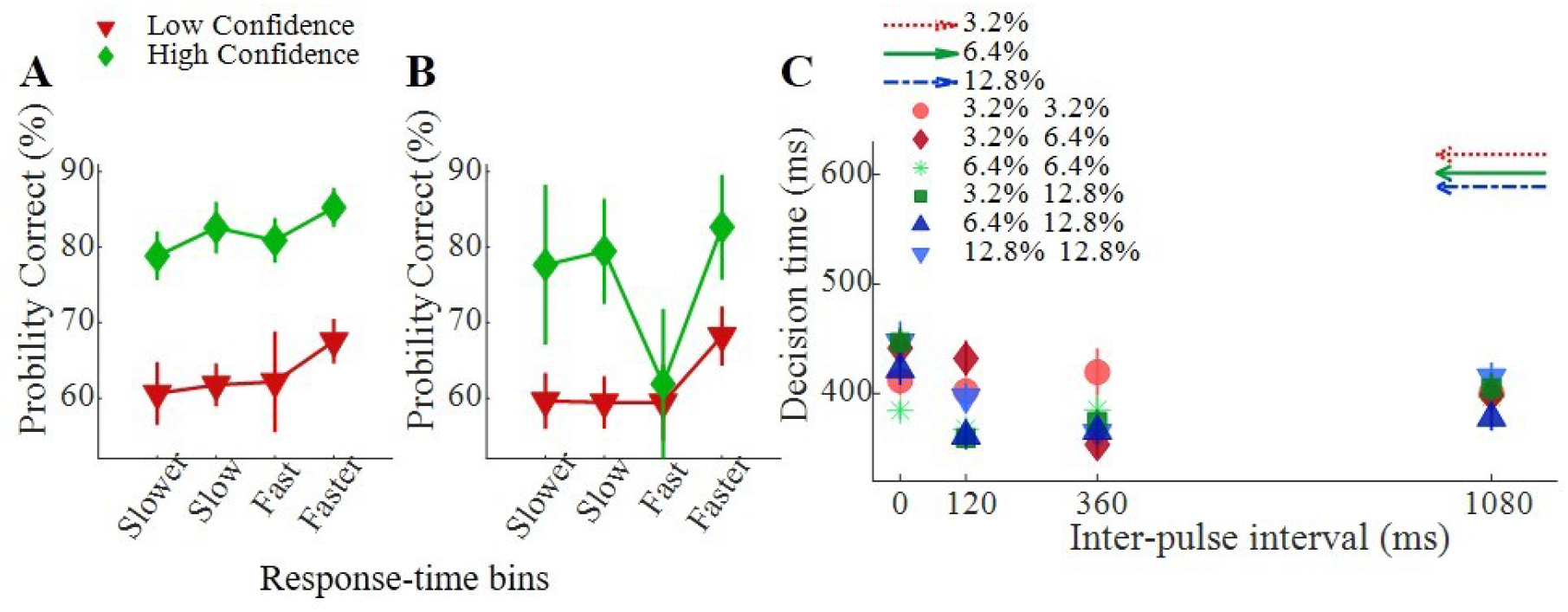
Response-time in single-pulse and double-pulse trials. **(A, B)** Accuracy as a function of response-time split by high (green) and low (red) confidence in double-pulse trials (A) and single-pulse trials (B). **(C)** Response-time of all coherence combinations clustered by interpulse intervals in double-pulse trials (dots) compared to single-pulse trials (arrows). In all panels, data are represented as group mean ± SEM.

### 3.6 EEG Analysis

We derived the ERPs of averaged signals for two confidence levels to verify whether there was a significant difference in the CPP across confidence levels. Figure 7 exhibits ERPs and scalp topographies for confidence levels time-locked to the stimulus onset in single-pulse trials. ERPs in the two confidence levels showed the ramping activities after 250 ms of the stimulus onset (Figure 7A). However, contrary to what we expected (Vafaei Shooshtari et al., 2019; Zizlsperger, Sauvigny, Händel, & Haarmeier, 2014), high confidence trials failed to confirm higher CPP amplitude in single-pulse trials. Accordingly, while, in single-pulse trials with low coherence (3.2% and 6.4%), there was no significant difference between CPP amplitude for low and high confidence trials (Figure 7C, 7D), there was a significant difference in 12.8% coherence (Figure 7B).

**Figure 7.**
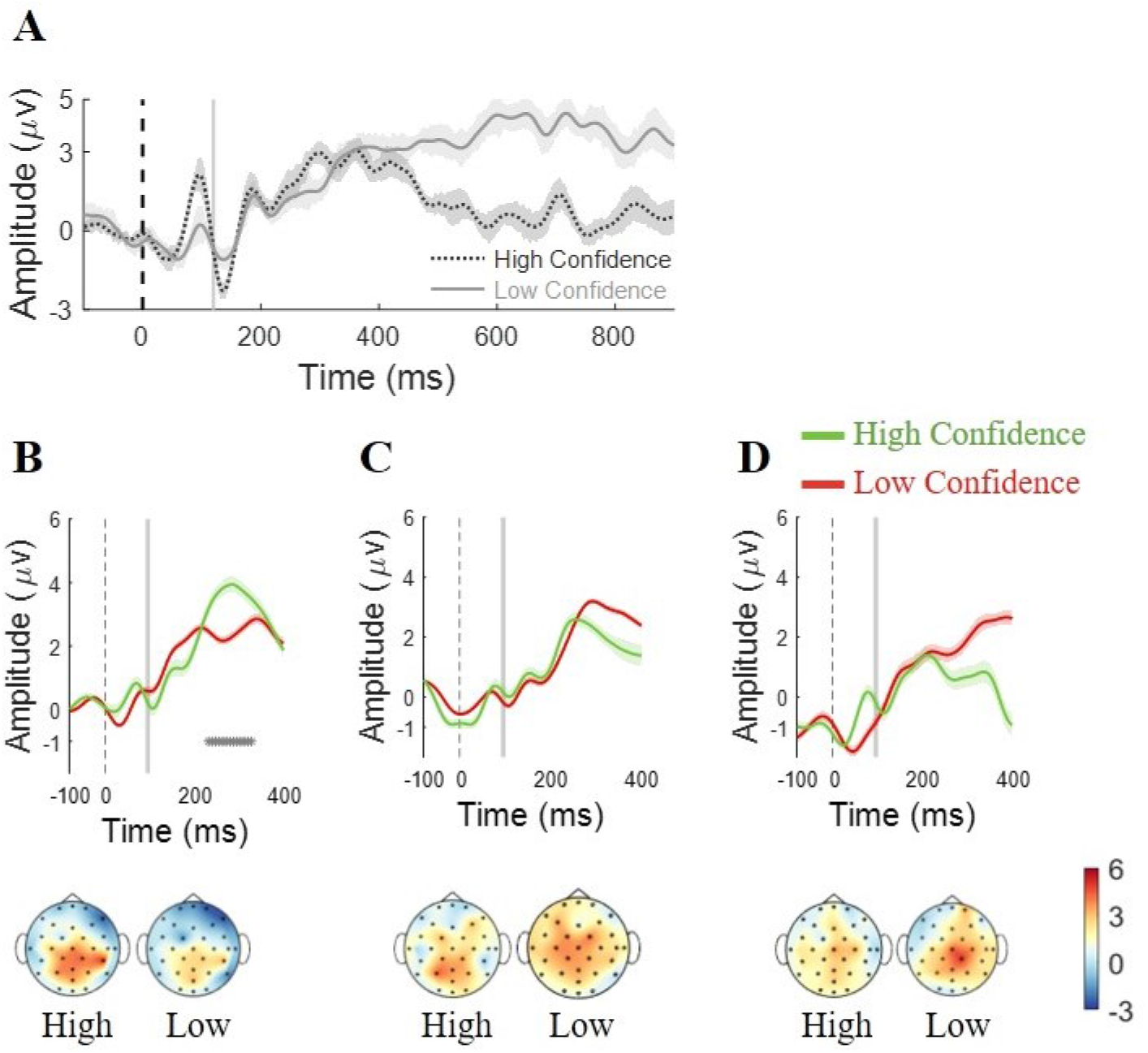
ERPs and scalp topographies in single-pulse trials. **(A)** ERP over the entire trial grouped by high and low confidence. ERPs and Scalp topographies in the two levels of confidence of single-pulse trials show significant difference in **(B)** 12.8% coherence, and insignificant difference in **(C)** 6.4% coherence, and **(D)** 3.2% coherence. Scalp topographies are represented in two levels of confidence (the mean amplitude in a time-window ranging from 200 ms to 500 ms after stimulus onset of second pulse). The shading region around the mean indicates SEM. * indicate p<.05 from a cluster-corrected *t*-test, of the difference between the two-time.

We also derived the ERPs of averaged signals for two confidence levels in double-pulse trials. ERPs in the two confidence levels showed the ramping activities after 250 ms of the stimulus onset and as we expected, high confidence trials confirmed higher CPP amplitude in double-pulse trials for all different interpulse intervals (see Figure 8A, 8B, 8C, and 8D over the entire trial). Moreover, Figure 8E, 8F, and 8G exhibit significant differences in CPP amplitude for two confidence levels time-locked to the stimulus onset in each pulse of double-pulse trials. As literature has confirmed that CPP amplitude is associated with confidence at the time of the response (Boldt et al., 2019), although ERPs for both pulses are presented, we only reported significancy for the second pulses. In adition, the previous findings distinguished the correlation between CPP signal and coherence (Kelly and O’Connell, 2013) and coherence is the decisive characteristic of confidence (Baranski and Petrusic, 1998; Van denBerg et al., 2016b). Accordingly, to verify that our findings are not relevant to different coherence, we only included double-pulse trials with equal motion strengths. Accordingly, ERPs and scalp topographies showed that even in trials with low coherence of double-pulse trials, there were significant difference between CPP amplitude for low and high confidence trials.

**Figure 8.**
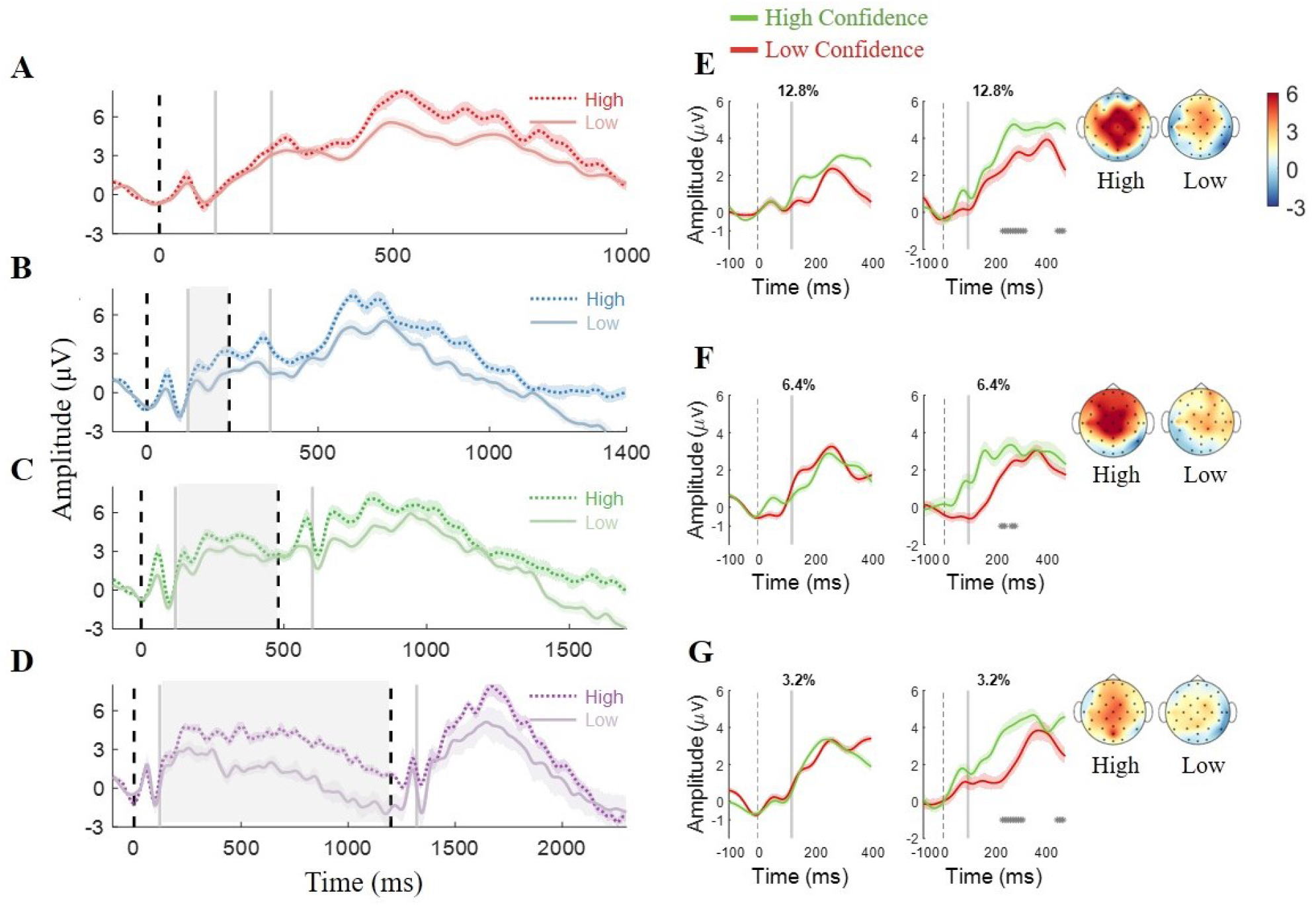
ERPs and scalp topographies in double-pulse trials. (**A-D**) ERPs over the entire trial grouped by high and low confidence in trials with zero, 120 ms, 360 ms, and 1080 ms interpulse intervals. **(E-G)** ERPs and Scalp topographies in the two levels of confidence are distinct after the stimulus onset. Motion strength are indicated at the top of each panel: 12.8%, 6.4%, and 3.2%. Scalp topographies are represented in two levels of confidence (the mean amplitude in a time-window ranging from 200 ms to 500 ms after stimulus onset of second pulse). The shading region around the mean indicates SEM. * indicate p<.05 from a t-test, of the difference between the two-time.

**Figure 9.**
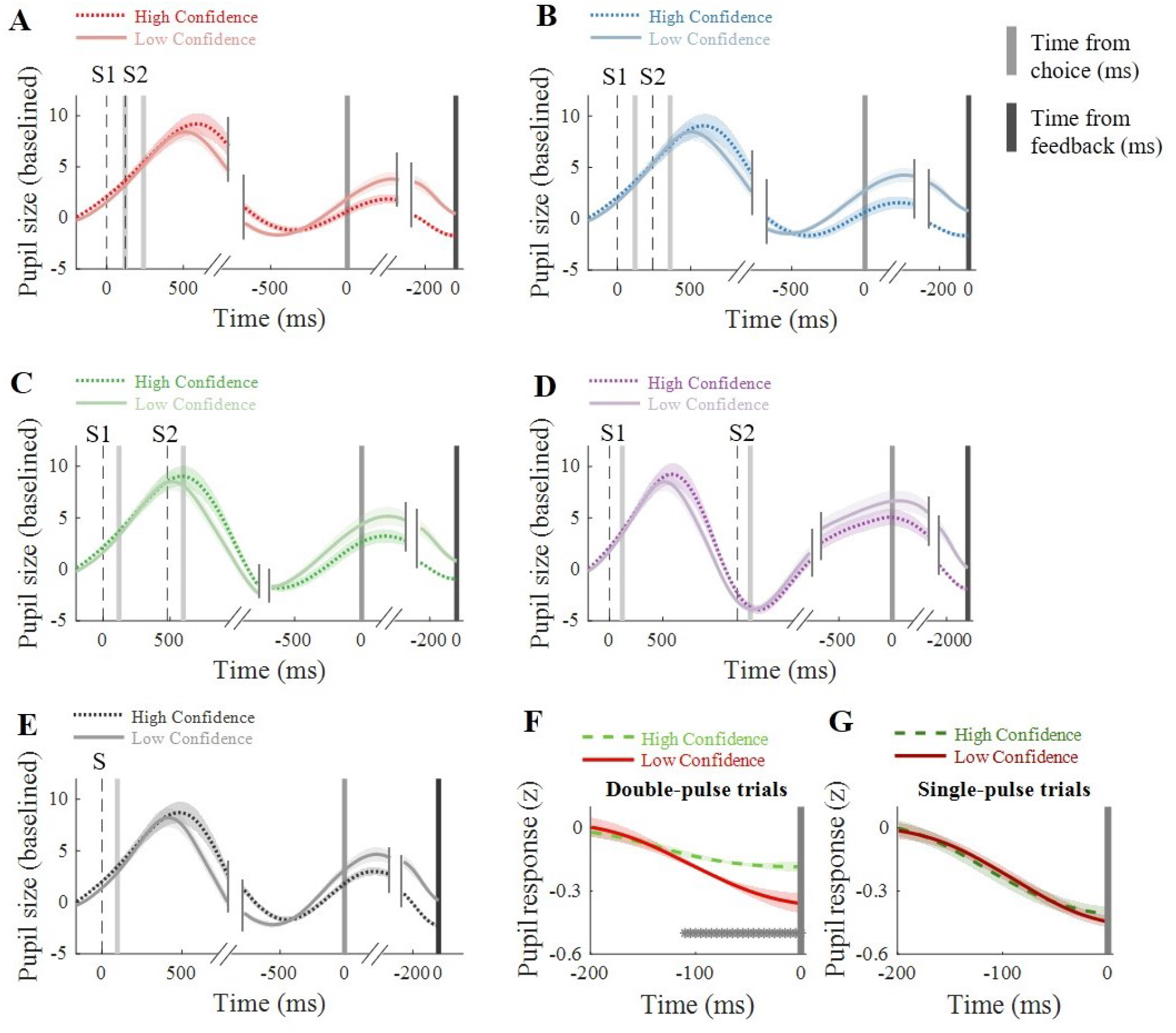
Pupil responses in single-pulse and double-pulse trials. **(A-D)** Time course of pupil responses throughout the entire trial grouped by high and low confidence in trials with zero, 120 ms, 360 ms, and 1080 ms interpulse intervals. **(E)** Time course of pupil responses throughout the entire trial grouped by high and low confidence in single-pulse trials. **(F, G)** Standardized pupil response across time-window aligned to the feedback, high confidence trials (green) vs. low confidence trials (red) in double-pulse trials and single-pulse trials. The shading region around the mean indicates SEM. * indicate p < .05 from a *t*-test in F and G.

Furthermore, a linear regression model confirmed the influence of confidence on CPP amplitude in double-pulse trials (*β_1_* = .88, *p* < .05, 95% CI = [.39, 2.20]; *f^2^* = .10), whereas there was not sufficient evidence to show the influence of confidence on CPP amplitude in single-pulse trials (*β_1_* = −6.58, *p* = .10, 95% CI = [−13.40, .24]; *f^2^* = .28). Additionally, a statistical comparison of the single-pulse and double-pulse trials represented that the difference of the CPP amplitude for high and low confidence responses depended on the type of trials (Eq.13, *β_3_* = 7.46, *p_bonf*3_* < .05, 95% CI = [3.09, 11.83], *f^2^* = .25).

To certify that our findings are not relevant to participant performance, we only analyzed the ERP signals in the two confidence levels only for correct responses. Supplementary Figure 11 expresses that the ERPs of the two confidence levels were independent of accuracy coding in the centro-parietal area, and they were associated with a degree of certainty.

### 3.7 Pupil responses

As shown in Figure 10A, 10B, 10C, 10D, and 10E, during decision formation in both single-pulse and double-pulse trials, pupil dilation in response to the confidence levels was not clearly distinct after the stimulus onset, this is in line with previous works (Vafaei Shooshtari et al., 2019). However, pupil responses were peaking immediately after making the choice, similar to the previous findings (Urai et al., 2017).

Additionally, to quantify the temporal evolution of confidence in the pupil, we took the mean baseline-corrected pupil signal during 200 ms before feedback delivery as our measure of pupil response. In line with previous work (Colizoli, De Gee, Urai, & Donner, 2018; Urai et al., 2017) pupil responses reflected decision confidence in our double-pulse trials (Figure 10F; *β_1_* = −.15, *p* < .001, 95% CI = [−.25, −.05], *f^2^* = .28), whereas in single-pulse trials, pupil responses did not reflect decision confidence (Figure 10G; *β_1_* = −.12, *p* = .21, 95% CI = [−.21, −.03], *f^2^* = .20). Furthermore, a statistical comparison of the single-pulse and double-pulse trials represented that the difference of the pupil response for high and low confidence responses depended on the type of trials (Eq.14, *β_3_* =.33, *p_bonf*3_* < .05, 95% CI = [−.09, .76], *f^2^* = .23). To certify that our findings are not relevant to participant performance, we only analyzed the pupil response signals in the two confidence levels for correct responses. Supplementary Figure 12 expresses that the pupil responses during 200 ms before feedback delivery of the two confidence levels were independent of accuracy, and they were associated with confidence in double-pulse trials.

## 4 Discussion

The current study was designed to clarify the confidence of decisions in a more similar context to the real world, where the evidence arrives separately. Using an experimental design adopting either single or double pulse(s) of RDM stimuli, in two separate studies, we integrated behavioral, EEG responses, and pupillometry to examine how people combined separate pieces of information to form their decision. We were particularly interested in how accuracy and confidence are related to each other in the context of decision-making. We predicted that similar to accuracy, confidence would improve when people integrate separate pieces of information, confidence would be leveraged larger by more recent information, and that using behavioral, neural, and pupillometry markers of confidence would be distinguishable in both single-pulse trials and double-pulse trials. In summary, we found that although participants’ confidence was invariant primarily to the interpulse interval, confidence scoring was not noticeably enhanced in double-pulse trials compared to single-pulse trials. Instead, participants only reported their confidence with higher resolution suggesting that adding an extra pack of information only increases participants’ metacognitive sensitivity. Moreover, participants used both pulses to decide about their confidence irrespective of their recency or primacy orders. Results also revealed that using RT, EEG, and pupillometry analysis allows a conceivable track of the confidence in double-pulse trials but not in single-pulse trials. Findings are discussed separately for behavioral, computational, and implicit confidence markers in the following subsections.

### 4.1 Behavioral and motion energy findings

Contrary to our hypothesis, and unlike accuracy, confidence ratings in double-pulse trials did not increase compared to single-pulse trials. In addition, our data failed to support our prediction that participants mainly trust the evidence of one of the pulses and ignore the other one. The trusted pulse can either be the first or second pulse; it also can simply be the stronger pulse. Nevertheless, the effect of sequence and interaction of pulses on confidence was examined and no effect was observed. Inconsistent with the previous claims that participants obtained more information from a second pulse (Kiani et al., 2013; Tohidi-Moghaddam et al., 2019), our motion energy analysis did not confirm the asymmetric influences of the two pulses for confidence. Yet our findings are in line with a large body of evidence (De Gardelle & Mamassian, 2015; Herce Castañón et al., 2019; Rahnev & Denison, 2018; Zylberberg et al., 2016, 2014) that shows that the observers do not make their decisions precisely following the confidence rating. Indeed, comparing confidence in double-pulse trials grouped by accuracy, we found that the participants had lower confidence in double-pulse trials than single-pulse trials for incorrect choices but higher confidence for correct choices. In other words, compared with single-pulse trials, in double-pulse trials, participants adjusted their confidence by enhancing their confidence resolution or metacognitive sensitivity. Typically, confidence facilitates evidence accumulation and drives a confirmation bias in perceptual decision-making (Rollwage et al., 2020). Likewise, we suggest that an extra brief and weak evidence can validate confidence and improve metacognitive sensitivity.

### 4.2 Computational modeling findings

To understand the nature of the differences of metacognitive sensitivity in double-pulse vs. single-pulse trials, we compared corresponding estimated metacognitive parameters for each participant. Likewise, we included the expected parameters that would be achieved in double-pulse trials under the assumption of perfect integration. Accordingly, we computed *Meta*–*d*’/*d*′ as a measure of ‘metacognitive efficiency’. In the case of *Meta*–*d*’ = *d*’, the observer is metacognitively ‘ideal’. Indeed, all the information available for the decision would be used to report the confidence. Yet, in many cases, we might find that *Meta*–*d*’ < *d*′, along with some degree of noise or suboptimality (Fleming & Lau, 2014; Maniscalco & Lau, 2012). Conversely, we may find that *Meta*–*d*’ > *d*′, if participants are able to draw on additional information such as hunches (Rausch & Zehetleitner, 2016), further processing of stimulus information (Charles, Van Opstal, Marti, & Dehaene, 2013) or extra prior knowledge on the task. In the model fitted to double-pulse trials, *Meta*–*d*’/*d*′ was around .8 and near to ideal for almost all participants. However, as it varied considerably between participants in single-pulse trials, the value could not be adjusted in the perfect integrator model too.

Previously, the better-than-expected performance in double-pulse trials was explained by underperformance in single-pulse trials (Kiani et al., 2013). In current study, metacognitive sensitivity in double-pulse trials surpasses the value predicted by the perfect integrator model (Figure 5C and Supplementary Figure 4). This effect can be followed in all of our participants (except one of the participants from Experiment 2) and can be explained by low confidence resolution in single-pulse trials.

Metacognitive noise is the noise that affects confidence estimates but not perceptual decisions (De Martino, Fleming, Garrett, & Dolan, 2013; Jang, Wallsten, & Huber, 2012; Maniscalco & Lau, 2016; Mueller & Weidemann, 2008; Rahnev, Nee, Riddle, Larson, & D’Esposito, 2016; Shekhar & Rahnev, 2018; Van den Berg, Yoo, & Ma, 2017). A recent work categorized sources of metacognitive inefficiency (Shekhar & Rahnev, 2020). Accordingly, metacognitive noise is a superordinate term for all noise sources that impact the confidence formation process (Shekhar & Rahnev, 2020, 2021), ranging from systematic to nonsystematic input and computation. Nevertheless, the exact source of metacognitive noise remains unclear (Shekhar & Rahnev, 2020). This noise can be traced in our perfect integrator model, which was capable of accumulating decision evidence perfectly but could not predict confidence formation in our task. We suggest that the perfect integrator model was unable to adjust to confidence criteria when predicting confidence in double-pulse trials. However, an improved SDT capable of addressing metacognitive noise might be able to empower the employed perfect integrator model. Furthermore, SDT is not the only available model to implement a perfect integrator model. Previous studies suggested attractor models as a candidate model to implement the perfect integrator model (Kiani et al., 2013; Waskom & Kiani, 2018). Attractor models are a group of networks that form a bridge between cognitive theory and biological data which exploits inhibition to achieve to compete among alternatives (Wang, 2002; Wong & Huk, 2008). Although these models can integrate momentary evidence to establish a decision, they have specific failure behaviors that would be apparent when gaps separate the sources of evidence in time (Kiani et al., 2013). Besides, when the stimulus is very short, mostly, none of the attractors can be reached, and the network would revert back to the resting state after the stimulus offset (Wang, 2002). Therefore, the choice would be assigned randomly. However, our experiments’ data represent a noteworthy performance in single-pulse trials, not supporting this expectation. Consequently, to implement a perfect integrator model by implementing an attractor model, a mechanism for simulating a very short stimulus might be considered. Moreover, our behavioral assays highlighted different relationships between confidence and accuracy in the different conditions of the task. So, a dedicated neural module with a plausible circuit of confidence might be a better option to implement a perfect integrator model. Recently, multi-layer recurrent network models have been developed to account for decision confidence mechanisms (Atiya, Rañó, Prasad, & Wong-Lin, 2019; Paz, Insabato, Zylberberg, Deco, & Sigman, 2016). These models consist of multiple layers of neural integrators and align with neural evidence of decision confidence (Kepecs, Uchida, Zariwala, & Mainen, 2008; Murphy, Robertson, Harty, & O’Connell, 2015), they are suggested to justify the observed behavior.

Furthermore, perceptual decisions are often modeled using ideal observers (e.g., SDT). However, a source of suboptimal behavior in decision-making is ‘lapse’ (Gold & Ding, 2013; Pisupati, Chartarifsky-Lynn, Khanal, & Churchland, 2021). The lapses are an additional constant rate of errors independent of the evidence strength (Gold & Ding, 2013; Pisupati et al., 2021). Lapse rate has been shown to increase with higher perceptual uncertainty (Pisupati et al., 2021) and would be accounted for by fitting extra parameters to psychometrics models. Accordingly, as the perfect integrator model was based on SDT, ignoring lapse in the single-pulse trials might lead to mis-estimation of decision parameters in double-pulse trials. Consequently, further models including the lapse parameters (Pisupati et al., 2021), may improve the perfect integrator model’s predictivity.

### 4.3 Implicit confidence markers

Although research suggests faster decisions accompanied by higher confidence (Kiani et al., 2014; Vafaei Shooshtari et al., 2019; van den Berg et al., 2016; Zylberberg et al., 2016), our results do not show such an association in the presence of a brief piece of evidence. Moreover, our participants decided much faster in double-pulse trials compared single-pulse trials. Therefore, we proposed that the decrease of response-time in double-pulse trials would be reflected with higher internal confidence. However, another hypothesis of this variation pointed to the extra time duration in double-pulse trials, which can be used to increase readiness to decide. Therefore, to explore the hypothesis, we regressed the delay-time before response cue onset and response-time in both single-pulse and double-pulse trials. If the variation of response-time was primarily dependent on extra delay time, the delay time should have had a considerable effect on response-time, especially in our 120ms single-pulse trials when the stimulus duration was concise and the delay time varied. Nevertheless, the effects in both double-pulse and single-pulse trials are weak. Accordingly, the hypothesis that faster decisions reflect higher confidence in double-pulse trials is supported. In addition, response-time in double-pulse trials traces the confidence unlike in single-pulse trials.

Our findings furthermore suggest that reported confidence might not follow the confidence marker in EEG response. We focused on the CPP —a neural correlate of perceptual processing believed to reflect evidence accumulation and correlated to confidence (Boldt et al., 2019; Herding et al., 2019; Rausch et al., 2020; Vafaei Shooshtari et al., 2019; Zizlsperger et al., 2014). However, our findings suggest that in the presence of a brief and weak stimulus, entirely unlike in double-pulse trials, CPP amplitudes show no significant variation in a high and low level of confidence. As confidence in single and double-pulse trials did not vary significantly, we suggest that variation of CPP amplitude shares more commonalities with implicit confidence measures rather than explicit confidence measures like ratings. Moreover, we propose that pupil response relation to confidence rating varies as the task condition changes; when participants access brief and weak stimuli, no association is detected, unlike in the presence of a pair of separated stimuli. Our current observations are not easily reconciled with existing theoretical accounts of the impact of the confidence level on pupil response (Allen et al., 2016; Lempert et al., 2015; Urai et al., 2017). Therefore, our study has thus theoretical implications for the confidence and decision formation and provides a new insight into the relationship between confidence and accuracy in perceptual decision making. In line with the theoretical account, the findings of this study can clear up some of the ambiguities of real-world decision-making when the evidence is separated in time. Considering the fact that it would be more probable that people change their mind when decisions are accompanied by lower confidence (Fleming, Putten, & Daw, 2018; Resulaj, Kiani, Wolpert, & Shadlen, 2009), knowing how to make decisions in a discrete environment and confidence, can help encourage more sustainable choices in marketing and behavioral economics. Also, people can integrate information from themselves and others based on their confidence (Bahrami, B. et al., 2010; Bang, D. et al., 2017). Accordingly, confidence can affect the extent of people pursue others or be persuaded by them (Petty, Briñol, & Tormala, 2002). To sum up, when participants access brief and mainly weak stimuli, the confidence ratings are not reliable, and confidence could not be traced from response-time, pupil, and EEG response. In other words, implicit confidence markers, in some cases, might be incapable of following the conscious confidence rating. This is in line with innovative findings abstracting implicit confidence measures from explicit confidence measures (Logan & Crump, 2010).

### 4.4 Limitations and future directions

Our results were grounded in assumptions of integration strategy in decision-making. However, this insight has recently been reconsidered (Carland, Marcos, Thura, & Cisek, 2016; Stine et al., 2020). An urgency-gate model might explain participants’ decisions better (Evans, Hawkins, Boehm, Wagenmakers, & Brown, 2017; Thura, Beauregard-Racine, Fradet, & Cisek, 2012) rather than an integration strategy such as perfect integration. A participant’s strategy could be something between no integration and perfect integration or in a completely different space of models (Stine et al., 2020) and might be changed depending on the task paradigm or even the participant’s internal state (Evans & Hawkins, 2019; Najafi & Churchland, 2018; Tsetsos, Gao, McClelland, & Usher, 2012). Consequently, further models to discuss the decision strategy in separated pulses could guide future works. In addition, future experiments could develop computational approaches and attempt to implement other scenarios in a discrete environment to study choice and confidence formation and examine the involved processes. In addition, although the vast number of trials for each participant allowed us to do a robust subject-wise analysis and our EEG study replicated the same behavioral and modeling data, our small sample size prevents us from making general claims. Nevertheless, future research might capitalize on our paradigm to provide a situation in which confidence remains persistent but metacognitive sensitivity improves. In this way, future research continues studying the neural basis of metacognitive ability and consciousness in addition to previous works (Feuerriegel, Blom, & Hogendoorn, 2021; Fleming & Dolan, 2012).

## 5 Conclusion

To sum, despite its limitations, the present study sheds new light on confidence formation, especially in perceptual decision-making when diverse temporal gaps separate a pair of visual cues. Our data suggest that accumulated evidence from both pulses shape confidence invariant to the interpulse intervals up to 1 s and the association of accuracy and confidence improved. However, the confidence would not follow the same characteristics as accuracy. For example, confidence would not be leveraged larger by more recent information. Moreover, we showed that the classic perfect integrator model merely highlighted evidence accumulation and ignored the effect of the metacognitive noise that affects confidence, so it failed to predict confidence. Finally, integrating evidence from two separated pieces of information supports the confidence to be traced from implicit markers like RT, EEG, and pupil responses, unlike the situation in which participants have to decide based on a brief and weak pulse of information.

## 7 Author Contributions

ZA: conceptualization, data acquisition, analysis, visualization, writing - original draft, writing - review and editing; SZ: conceptualization, supervision, writing - review and editing; AJ: supervision, writing - review and editing; RE: conceptualization, supervision, writing - review and editing.

## 8 Acknowledgments

Data were recorded in the Cognitive Science Laboratory of Shahid Rajaee University. The authors would like to thank Samuel Klein for their constructive comments and proofreading of the manuscript. We thank all our participant and finally, special thanks to the open science movement and generous researchers who we had their helpful comments during the implementation and analysis.

## 9 Ethics

The ethics committee of the Iran University of Medical Sciences (protocol #IR.IUMS.REC1399648) approved the experimental protocol, and participants gave written informed consent.

## 10 Supplementary Material

The Supplementary Material for this article can be found online at:

## 11 Data Availability Statement

The datasets generated for this study are available on request to the corresponding author.

## Supplementary Material

### 13 Supplementary Appendix 1: Figures and Tables

#### 13.1 Supplementary Tables

**Supplementary Table 1.** Subtraction of confidence in double-pulse trials from single-pulse trials was significantly affected by choice accuracy.

| Participant | $\beta_1$ |
| --- | --- |
| P<sub>1</sub> | 0.17 ($p < 0.01$)<br>CI = [.15, .19] |
| P<sub>2</sub> | 0.21 ($p < 0.01$)<br>CI = [.19, .23] |
| P<sub>3</sub> | 0.10 ($p < 0.01$)<br>CI = [.08, .12] |
| P<sub>4</sub> | 0.12 ($p < 0.01$)<br>CI = [.10, .14] |

Each row shows the coefficients of Equation 5, the corresponding $p$ values, and 95% confidence interval.

**Supplementary Table 2.**
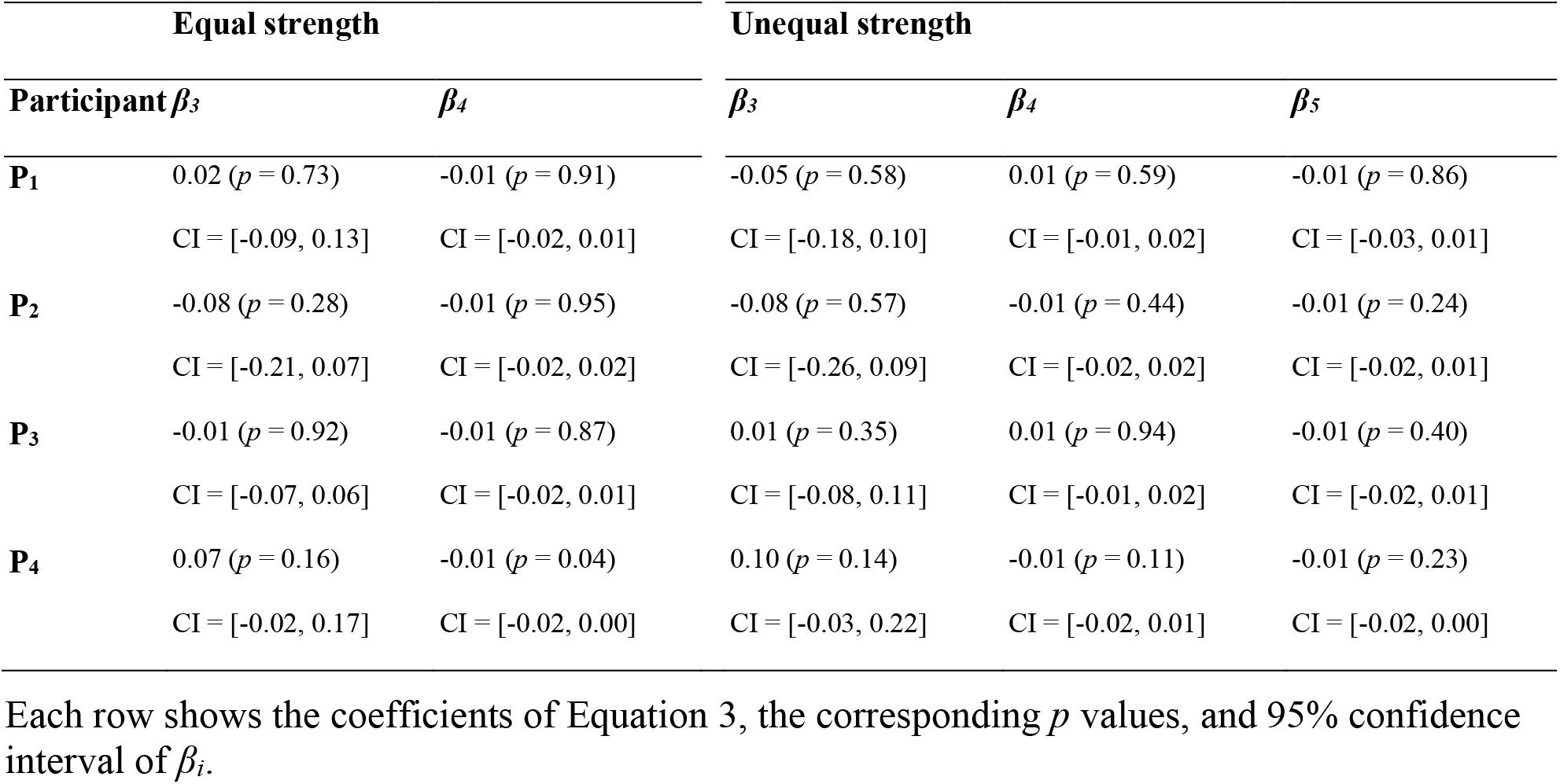
Performance was largely unaffected by interpulse interval for double-pulse trials with equal pulse strength and unequal pulse strength for each participant.

**Supplementary Table 3.** Performance was unaffected by interpulse interval for double-pulse trials with equal pulse strength and unequal pulse strength included trials with zero interpulse interval and one of the nonzero interpulse interval.

|  | Equal strength |  | Unequal strength |  |  |
| --- | --- | --- | --- | --- | --- |
| | $\beta_3$ | $\beta_4$ | $\beta_3$ | $\beta_4$ | $\beta_5$ |
| **0 vs. 120** | -0.14 ($p = 0.61$) | -0.03 ($p = 0.33$) | -0.45 ($p = 0.22$) | 0.03 ($p = 0.18$) | -0.01 ($p = 0.71$) |
|  | CI = [-0.69, 0.41] | CI = [-0.09, 0.32] | CI = [-1.19, 0.27] | CI = [-0.02, 0.09] | CI = [-0.06, 0.04] |
| **0 vs. 360** | 0.20 ($p = 0.08$) | 0.03 ($p = 0.04$) | -0.03 ($p = 0.79$) | 0.01 ($p = 0.83$) | -0.01 ($p = 0.13$) |
|  | CI = [0.01, 0.39] | CI = [0.01, 0.06] | CI = [-0.27, 0.21] | CI = [-0.02, 0.02] | CI = [-0.03, 0.01] |
| **0 vs. 1080** | 0.01 ($p = 0.91$) | -0.01 ($p = 0.21$) | -0.01 ($p = 0.73$) | 0.01 ($p = 0.98$) | -0.01 ($p = 0.58$) |
|  | CI = [-0.06, 0.07] | CI = [-0.02, 0.01] | CI = [-0.09, 0.07] | CI = [-0.01, 0.02] | CI = [-0.02, 0.01] |

Each row shows the coefficients of Equation 3, the corresponding $p$ values, and 95% confidence interval of $\beta_i$ .

**Supplementary Table 4.**
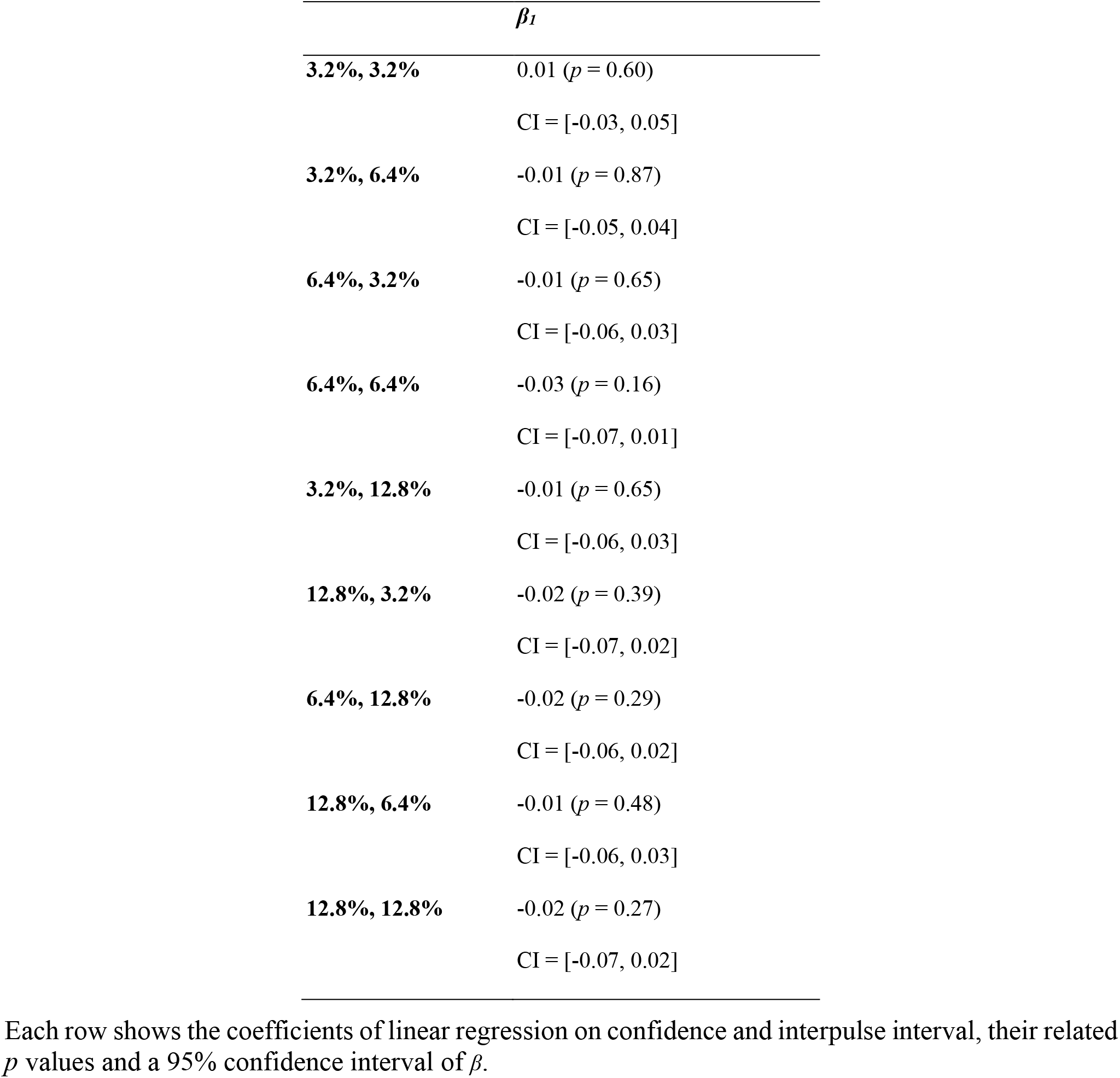
Confidence was unaffected by interpulse interval for double-pulse trials across all possible coherence sequence.

**Supplementary Table 5.**
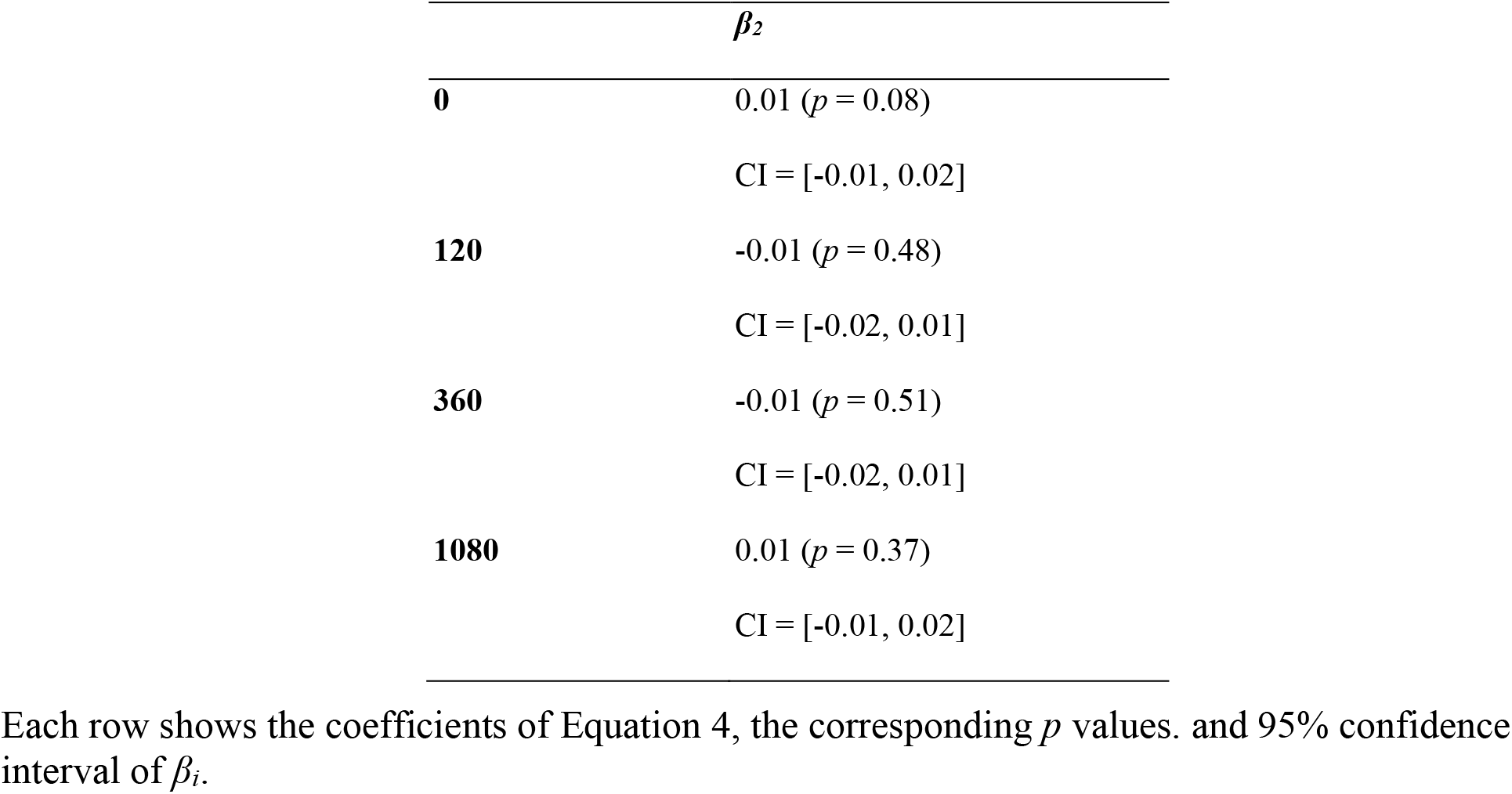
The order of the pulses failed to show significant effects on confidence on any of interpulse intervals.

**Supplementary Table 6.**
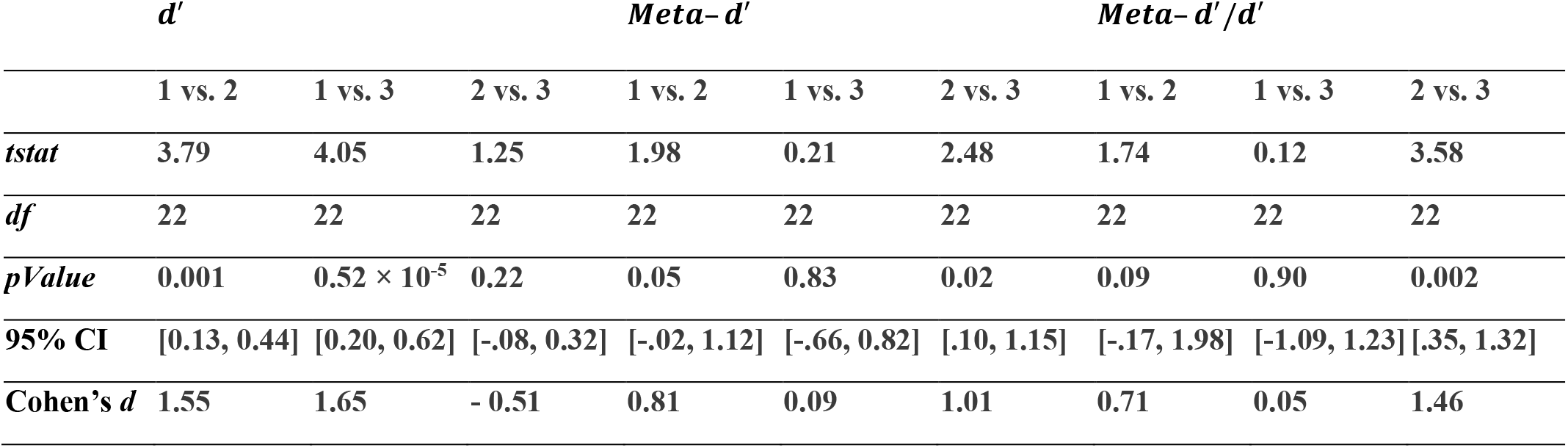
Pairwise comparisons across models (1: single-pulse trials, 2: double-pulse trials, 3: perfect integrator) for SDT parameters.

|  | *d'* |  |  | *Meta-d'* |  |  | *Meta-d' / d'* |  |  |
| --- | --- | --- | --- | --- | --- | --- | --- | --- | --- |
|  | 1 vs. 2 | 1 vs. 3 | 2 vs. 3 | 1 vs. 2 | 1 vs. 3 | 2 vs. 3 | 1 vs. 2 | 1 vs. 3 | 2 vs. 3 |
| *tstat* | 3.79 | 4.05 | 1.25 | 1.98 | 0.21 | 2.48 | 1.74 | 0.12 | 3.58 |
| *df* | 22 | 22 | 22 | 22 | 22 | 22 | 22 | 22 | 22 |
| *pValue* | 0.001 | $0.52 \times 10^{-5}$ | 0.22 | 0.05 | 0.83 | 0.02 | 0.09 | 0.90 | 0.002 |
| 95% CI | [0.13, 0.44] | [0.20, 0.62] | [-.08, 0.32] | [-.02, 1.12] | [-.66, 0.82] | [.10, 1.15] | [-.17, 1.98] | [-1.09, 1.23] | [.35, 1.32] |
| Cohen's *d* | 1.55 | 1.65 | - 0.51 | 0.81 | 0.09 | 1.01 | 0.71 | 0.05 | 1.46 |

**Supplementary Table 7.** ***d*′, *Meta*–*d*′, and *Meta*–*d*′/*d*′** was unaffected by interpulse interval for double-pulse trials in both experiments.

| | | $d'$ | $Meta-d'$ | $Meta-d'/d'$ |
| --- | --- | --- | --- | --- |
| **Experiment 1** | $\beta_i$ | -0.08 ($p = 0.33$) | -0.23 ($p = 0.09$) | -0.18 ($p = 0.16$) |
|  |  | CI = [-0.22, 0.07] | CI = [-0.48, 0.03] | CI = [-0.43, 0.07] |
| **Experiment 2** | $\beta_i$ | -0.07 ($p = 0.38$) | 0.09 ($p = 0.49$) | 0.15 ($p = 0.30$) |
|  |  | CI = [-0.23, 0.08] | CI = [-0.17, 0.35] | CI = [-0.13, 0.43] |

Each cell shows the coefficients of linear regression on SDT’s parameters and interpulse interval, the corresponding $p$ values and 95% confidence interval of $\beta_i$ .

#### 13.2 Supplementary Figures

**Supplementary Figure 1.**
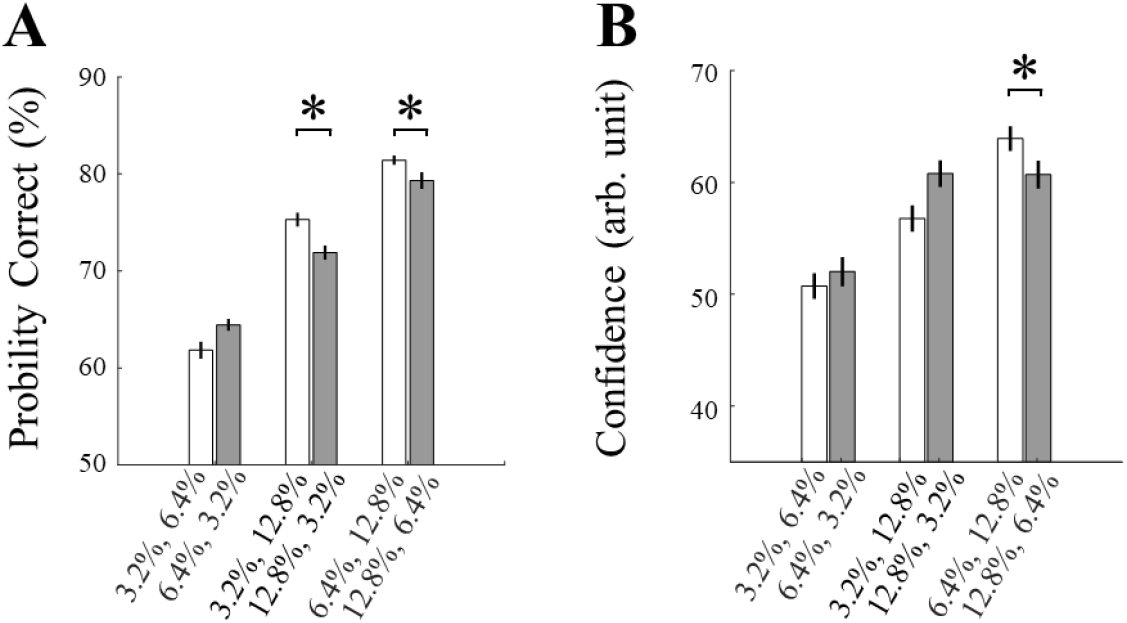
Choice confidence was not depended on the sequence of motion pulses in Experiment 2. **(A)** The weak–strong pulse sequence contributed higher accuracy than the strong–weak sequence. **(B)** The weak–strong pulse sequence did not contribute higher confidence compared to the strong–weak sequence. In all panels, data are represented as group mean ± SEM. (\**p*<0.05).

**Supplementary Figure 2.**
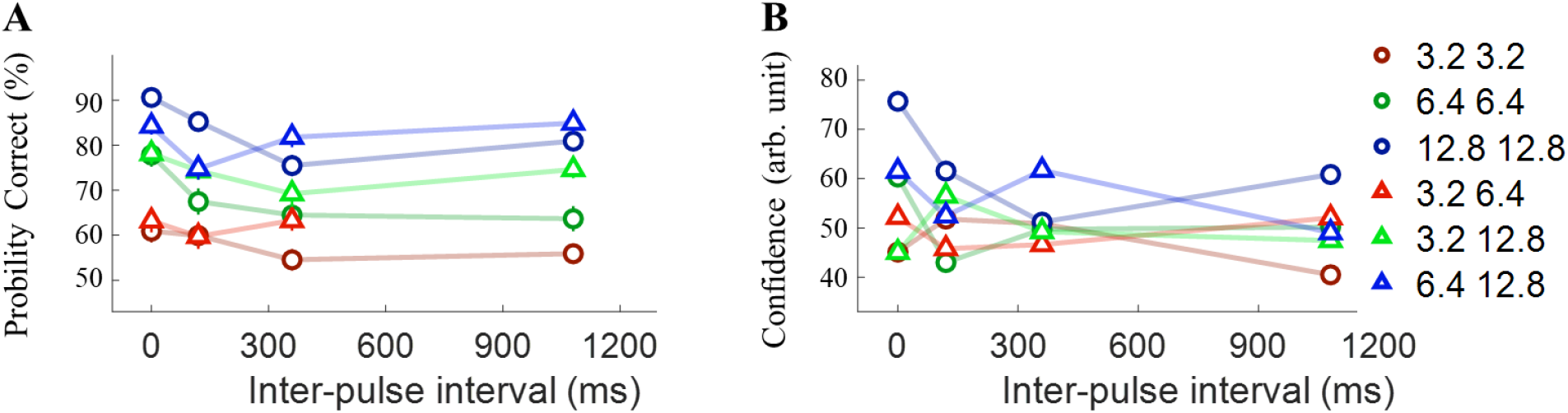
The Interplay between confidence, accuracy, and interpulse intervals in double-pulse trials in Experiment 2. **(A)** Choice accuracy for double-pulse trials grouping in all possible interval conditions. **(B)** Pooling data across all time intervals calculated the confidence of double-pulse trials. Each data point addresses pooled data from indicated sequence pulse and its reverse order (e.g., 12.8–3.2% and 3.2 −12.8%).

**Supplementary Figure 3.**
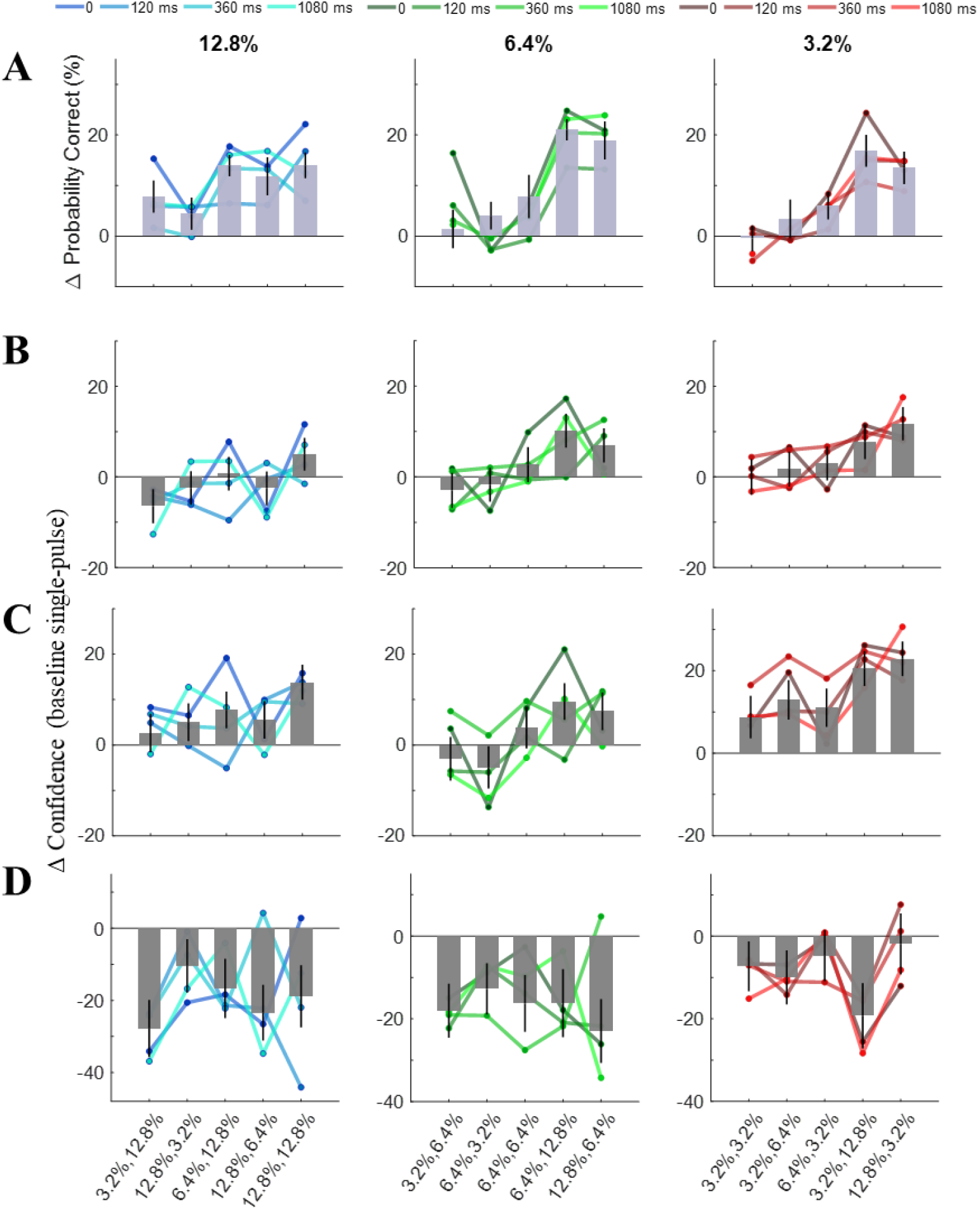
Variation of accuracy or confidence in double-pulse trials baselined by the corresponding coherence (3.2%, 6.4% and 12.8% for each column) in Experiment 2. **(A)** Considering all the trials, accuracy improved in combination with almost all pulses compared to the baseline. **(B)** Considering all the trials, confidence improved in combination with stronger pulses, whereas the confidence in sequence with a weaker pulse either decreased or remained constant. **(C)** In correct-choice trials, the increasing effect of stronger pulses is more obvious and the confidence even slightly improved in combination with weaker pulses compared to the corresponding baseline. **(D)** Interestingly, in incorrect trials, the confidence decreased in every condition. The colored line representing matching data for each of four possible gaps. In the bar graph, the data are represented as group mean ± SEM.

**Supplementary Figure 4.**
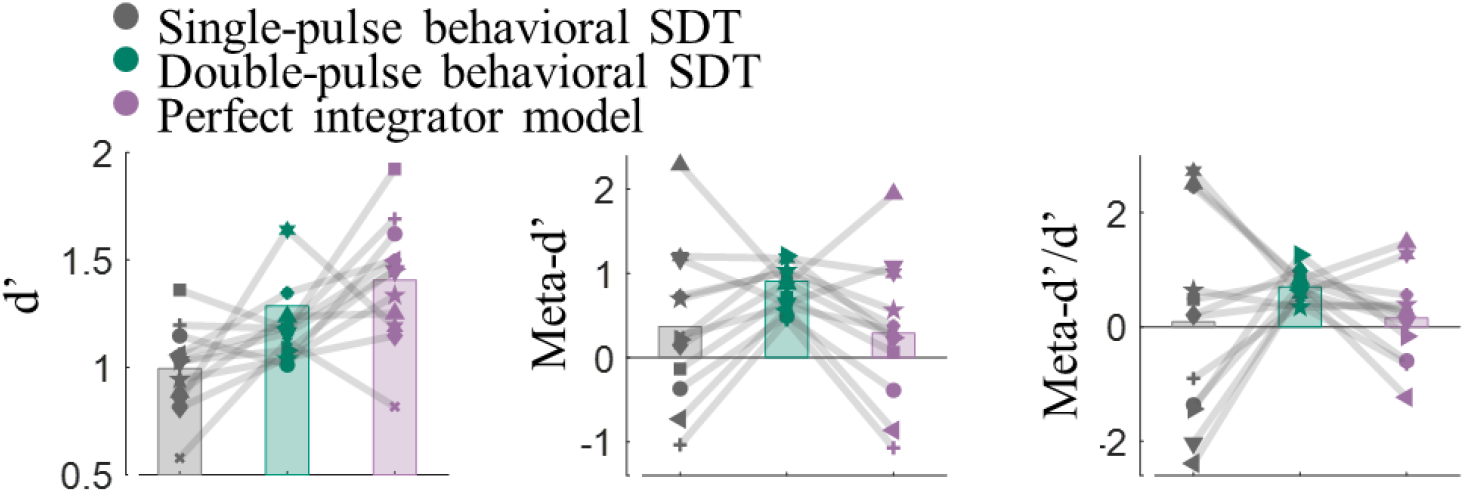
Comparison of models and human behavior in Experiment 2. Stimulus sensitivity (*d′*), metacognitive sensitivity (*Meta–d′*), and metacognitive efficiency (*Meta–d′/d′*) estimated for single-pulse trials, double-pulse trials and the perfect integrator models.

A univariate ANOVA showed that *d*′ between models fitted to double or single-pulse trials and the perfect integrator model significantly differed (F(2,33) = 9.99; *p* = 0.41 × 10^−4^). Also, a univariate ANOVA showed that *Meta*–*d*′ between models fitted to double-pulse or single-pulse trials and the perfect integrator model partially differed (F(2,33) = 1.04; *p* = 0.09). We also computed metacognitive efficiency (*Meta–d′/d′*), A univariate ANOVA revealed an insignificant difference on all three models (F(2,33) = 2.50; *p* = 0.10).), We also applied the *t*-test as a post hoc procedure to compare all pairs of *d*′, *Meta*–*d*′, *Meta*–*d*′/*d*′ calculated in three models (**Supplementary Table 5**).

**Supplementary Figure 5.**
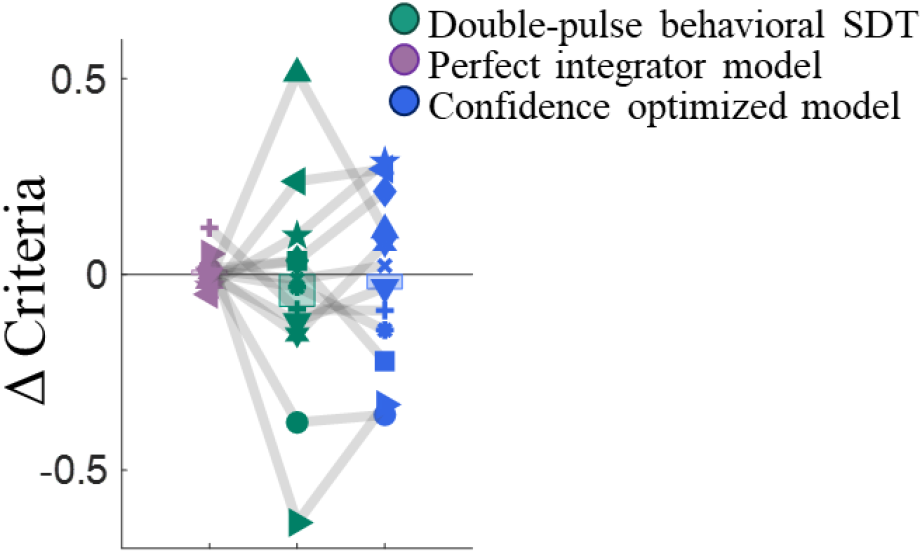
Variation of confidence criteria baselined to single-pulse trials for the perfect integrator model vs. the double-pulse trials model and the type 2 optimized model in Experiment 2.

**Supplementary Figure 6.**
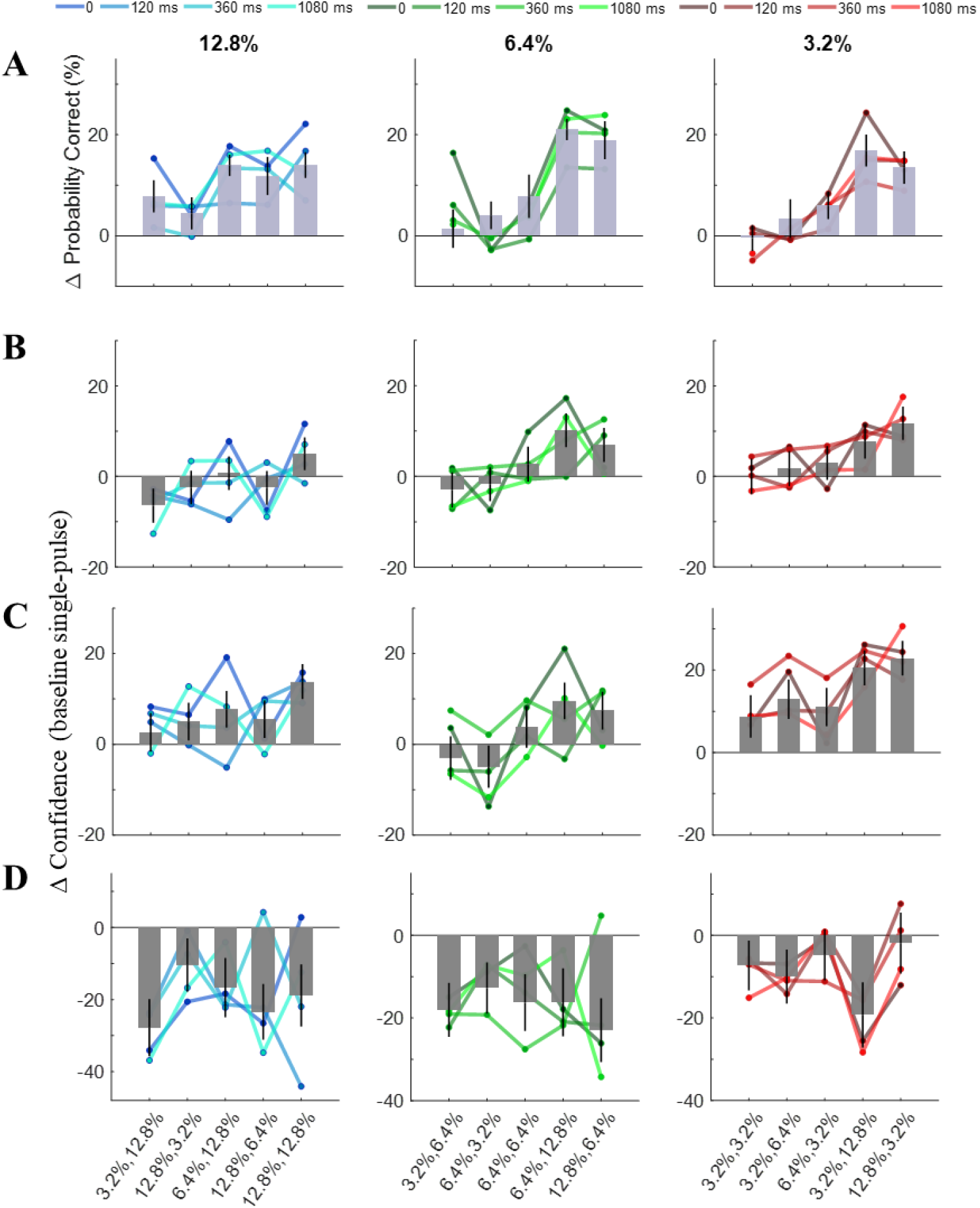
Variation of accuracy or confidence in double-pulse trials baselined by the corresponding coherence (3.2%, 6.4%, and 12.8% for each column) for the different interpulse intervals in Experiment 1. **(A)** Considering all the trials, accuracy improved in combination with almost all pulses compared to the baseline. **(B)** Considering all the trials, confidence improved in combination with stronger pulses, whereas the confidence in sequence with a weaker pulse either decreased or remained constant. **(C)** In correct-choice trials, the increasing effect of stronger pulses is more significant and the confidence even slightly improved in combination with weaker pulses compared to the corresponding baseline. **(D)** Interestingly, in incorrect trials, the confidence decreased in every condition—the colored line representing matching data for each of four possible gaps. In the bar graph, the data are represented as group mean ± SEM.

**Supplementary Figure 7.**
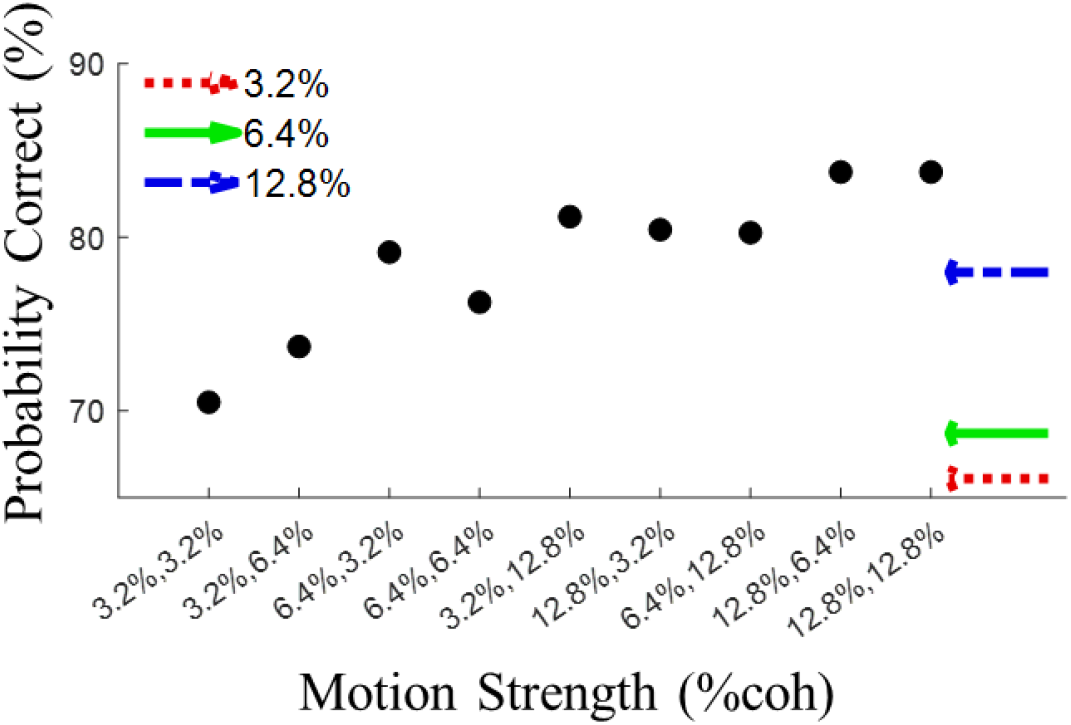
Accuracy in single and double pulse(s) trials when the same number of trials from each coherence with the same confidence range is selected randomly.

**Supplementary Figure 8.**
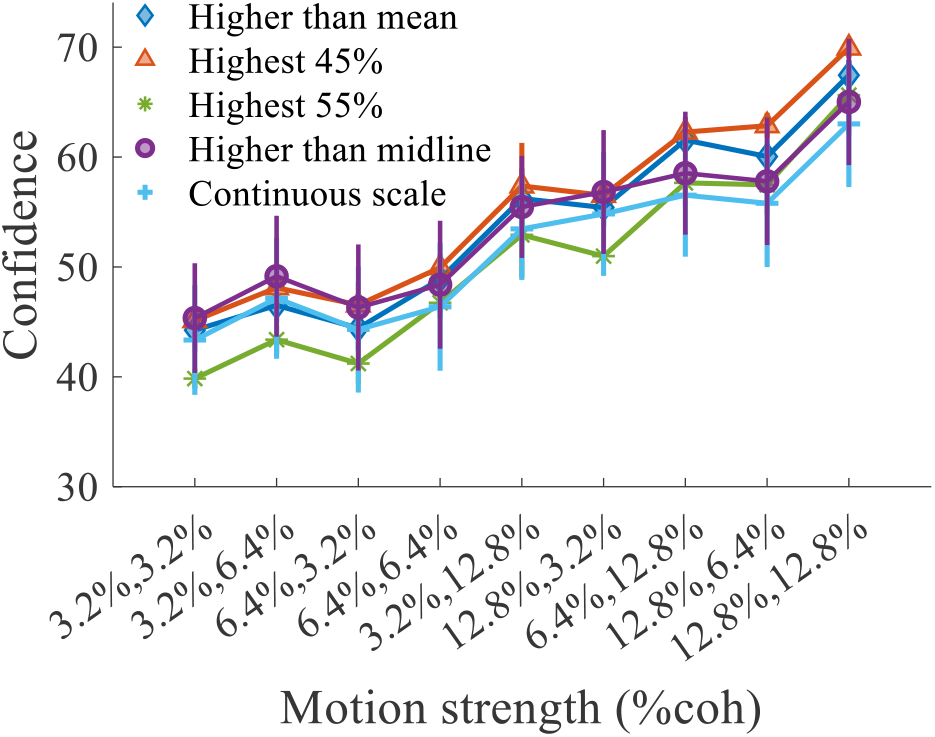
Graded confidence and four methods of confidence categorization. A univariant ANOVA showed that graded confidence and probability of high confidence calculated by four different approaches for each participant in double-pulse trials did not significantly differ (F(4,145) = 7.99; *p* = 0.41). Four paired-samples *t*-tests between the probability of high confidence calculated by different categorization methods and graded confidence showed no difference (all *ps* > 0.10).

**Supplementary Figure 9.**
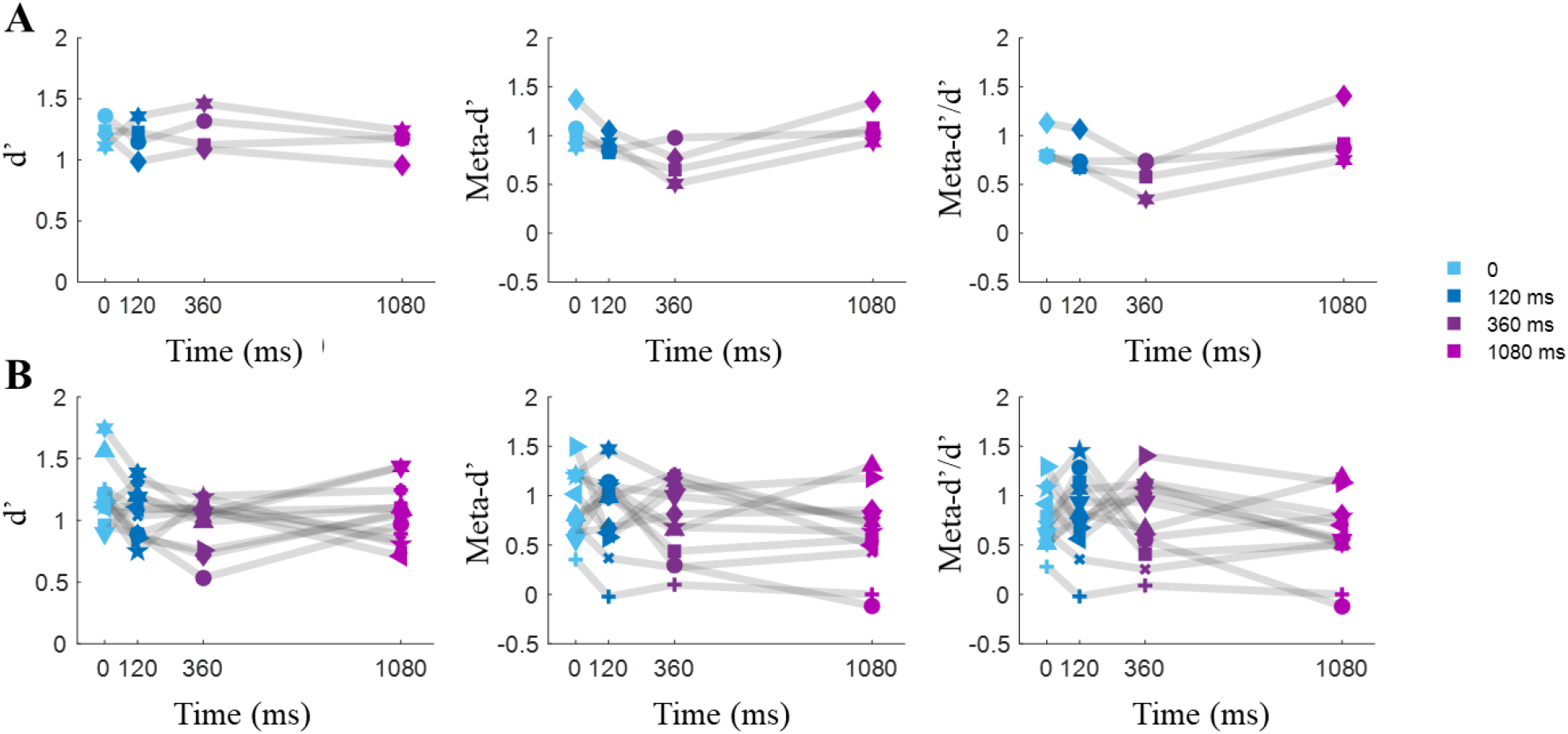
Stimulus sensitivity (*d′*), metacognitive sensitivity (*Meta–d′*) and, metacognitive efficiency (*Meta–d′/d′*) estimated for double-pulse trials were independent of the interpulse interval in **(A)** Experiment 1 and **(B)** Experiment 2. Each dot represents the data of each participant.

**Supplementary Figure 10.**
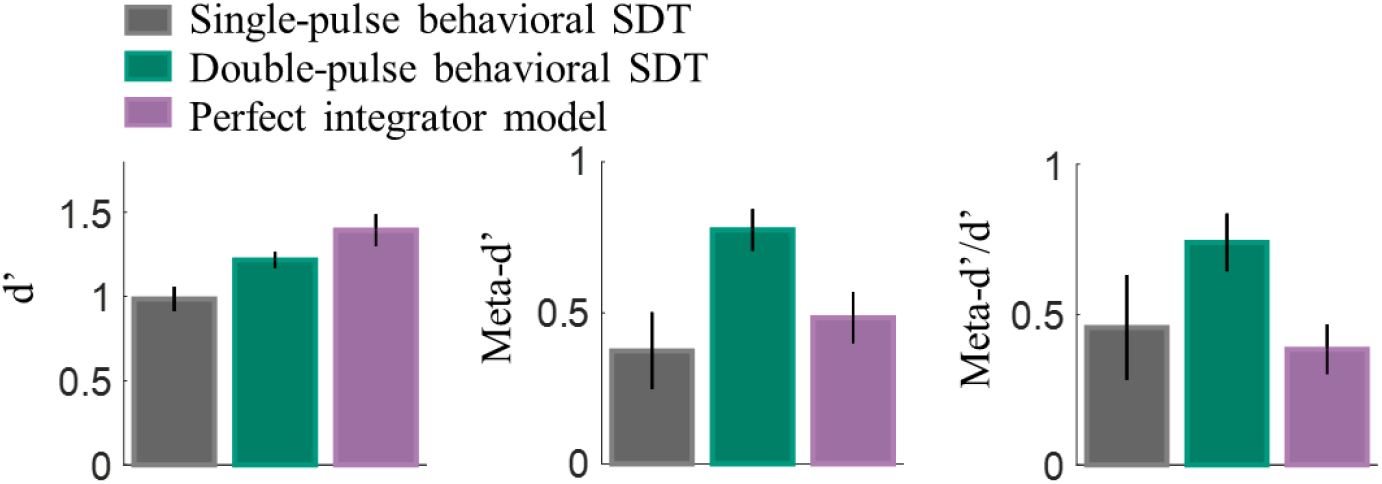
Comparison of models and human behavior considering the same numbers of trials in Experiment 1. Stimulus sensitivity (*d*′), metacognitive sensitivity (*Meta–d′*), and metacognitive efficiency (*Meta–d′/d′*) estimated for single-pulse trials, double-pulse trials, and the perfect integrator model.

We compared *d*′, *Meta– d*′ and, *Meta*–*d*′/*d*′ of fitted models to single/double-pulse trials and simulated data by the perfect integrator model, following up with three Dunn pair tests. A Kruskal-Wallis test showed that *d*′ between models fitted to double/single-pulse trials and the perfect integrator model did not significantly differ (*H*(3) = 3.23; *p* = 0.20). We also applied the Dunn test as a post hoc procedure to compare all pairs of *d*′ from three models. No *d*′ in models significantly differed from others (all *ps* > 0.21).

Also, a Kruskal-Wallis test showed that *Meta*–*d*′ between models fit to double/single-pulse trials and the perfect integrator model significantly differed (*H*(3) = 6.96; *p* = 0.03). Post-hoc Dunn was used to compare all pairs of *Meta*–*d*′ from three models. The difference of *Meta*–*d*′ of single-pulse and double-pulse trials was significant (*p* = 0.03, *CI* = [−12.58, −0.41]). However, the difference of *Meta*–*d*′ was insignificant for single-pulse trials and the perfect integrator model (*p* = 0.87, *CI* = [−7.83, 4.33]) and for double-pulse trials and the perfect integrator model (*p* = 0.17, *CI* = [4.75, 10.83]).

We also computed metacognitive efficiency (*Meta–d′/d′*). A Kruskal-Wallis test revealed a significant difference on all three models (*H*(3) = 7.42, *p* = 0.02). Metacognitive efficiency in double-pulse and the perfect integrator model differed significantly (*p* = 0.04, *CI* = [0.16 12.33]) while in double-pulse and single-pulse models, partially differed (*p* = 0.07, *CI* = [−11.83, 0.33]). The difference of single-pulse and the perfect integrator model was not significant (*p* = 0.99, *CI* = [−5.58 6.58]).

**Supplementary Figure 11.**
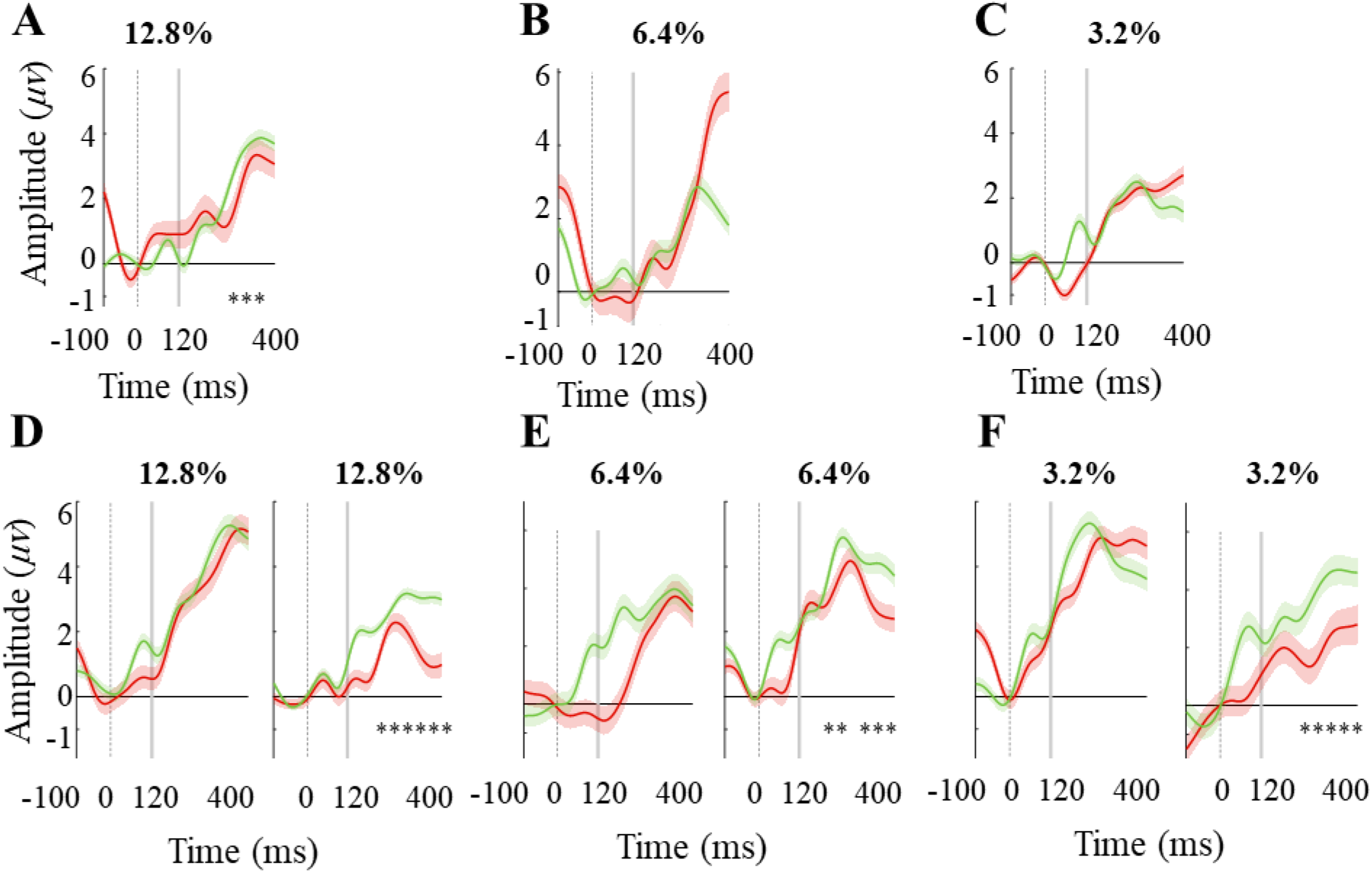
ERPs in single and double-pulse trials in correct trials. **(A) (B) (C)** ERPs of correct single-pulse trials shows an insignificant difference in weaker motion strength in high and low confidence level trials. **(D) (E) (F)** ERPs of correct trials in the two levels of confidence are distinct after the stimulus onset. The shading region around the mean indicates SEM. * indicate p<.05 from a *t*-test of the difference between the two-time.

**Supplementary Figure 12.**
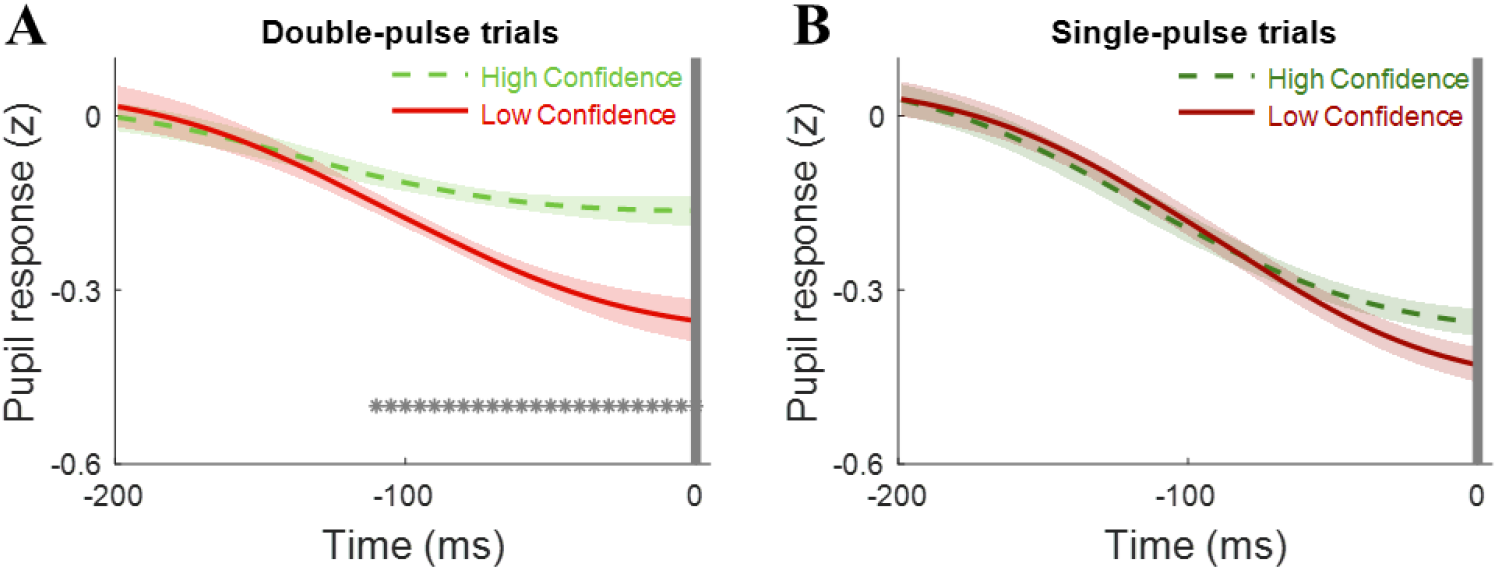
Standardized pupil response across time-window aligned to the feedback in correct trials, high confidence trials (green) vs. low confidence trials (red) in: **(A)** double-pulse trials, **(B)** single-pulse trials. The shading region around the mean indicates SEM. * indicate p < .05.

### 14 Supplementary Appendix 2: Signal detection theory models

In the binary decision, the observer must discriminate between stimuli labeled *S*_2_ or labeled *S*_1_. Each stimulus presentation generates a value on an internal decision axis (**Figure 1B**), corresponding to the evidence in favor of *S*_1_ or *S*_2_. Evidence generated by each stimulus class is normally distributed across the decision axis, and the distance between these distributions in standard deviation units (*d*′) which measures how well the observer can discriminate *S*_1_ from *S*_2_ (Maniscalco & Lau, 2012). The observer sets a decision criterion cr, such that all signals exceeding *cr* are labeled *S*_2_ and all those failing to exceed *cr* are labeled *S*_1_. The observer also sets criteria *cr*_2,“*S*1”_ and *cr*_2,“*S*2”_ to determine confidence ratings around the decision criterion cr. These two thresholds must be well-ordered so that *cr*_2,“*S*1”_ < *cr* < *cr*_2,“*S*2_ (**Figure 1B**). When a *S*_2_ response is made, a confident *S*_2_ response requires the evidence also to have surpassed the *cr*_2,“*S*2”_ threshold (Maniscalco & Lau, 2012, 2014).

#### 14.1 Confidence *Hit Rate* and *False Alarm Rate*

Sweeping the *cr*_2,“*S*2”_ criterion across the decision axis generates different values of confidence false alarm rate (*Prob(conf = “h”* | *stim* ≠ *resp*) and confidence hit rate (*Prob(conf = “h”* | *stim = resp*). A summary of the observer’s confidence performance is provided by hit rate (Hit Rate2) and false alarm rate (False Alarm Rate2) (Maniscalco & Lau, 2014):

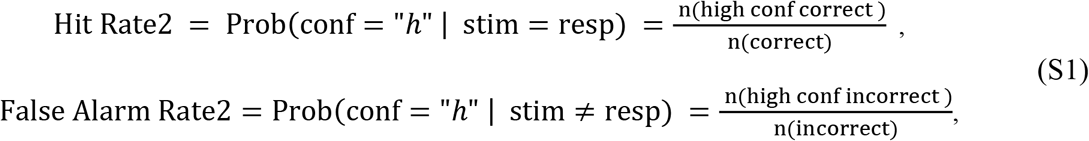

where *n*(*cond*) denotes a count of the total number of trials satisfying the condition *cond*.

#### 14.2 Decision *Hit Rate* and *False Alarm Rate*

In the SDT model, the decision hit rate (Hit Rate1) and the decision false alarm rate (False Alarm Rate1) are also calculated as follows:

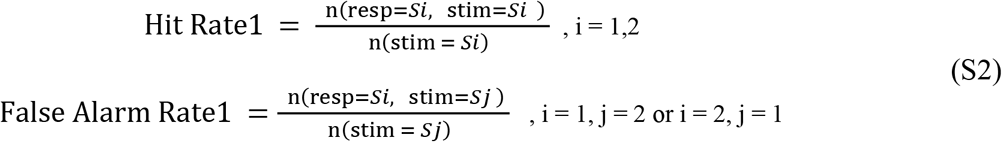

where *i* and *j* represent the stimulus classification. After calculating the Hit Rate1 and False Alarm Rate1 of each participant, *d*′ and *cr* are calculated as follows for each participant:

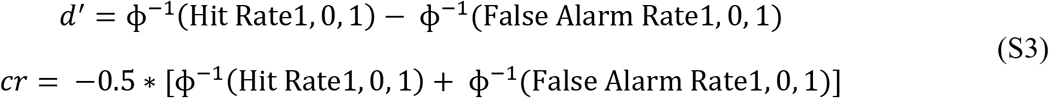

here, ϕ ^−1^ is the inverse of a function that represents a normal cumulative distribution and is calculated as follows:

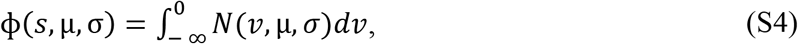

where *N*(*v*, μ, *σ*) is a Normal distribution with mean (μ) and standard deviation (*σ*). After the above computations, to simplify, we may consider the value of *cr* as zero point and move the distribution diagrams related to each option on the axis of the evidence.

By setting *d*′, *cr* and two criteria *cr*_2,“*S*1”_ and *cr*_2,“*S*2”_ (**Figure 1B**), the probabilities of each confidence rating conditional on a given stimulus and response (Hit Rate2 and False Alarm Rate2) can be calculated theoretically according to the following equations:

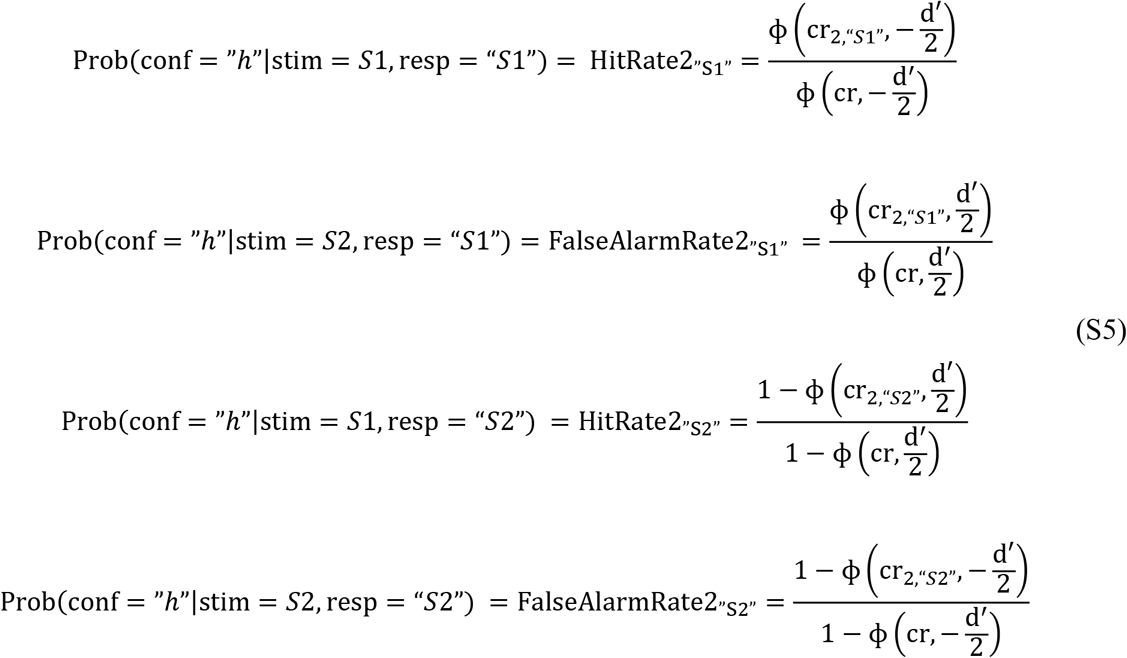

In the SDT model, there are different methods for adjusting the model with the data observed. In the method we used, *d*′ and *cr* were calculated from the participants’ performance (Eq. S3). Then, using maximum likelihood estimation (MLE) and Eq. S1 and S5 and by altering the value of the confidence criteria while holding *d*′ and *cr* constant, a set of (Hit Rate2, False Alarm Rate2) pairs ranging between (0, 0) and (1, 1) were generated. Moreover, *Meta*–*d*′ was found by fitting the decision SDT model to response-specific confidence.

